# Genomic landscape of the medieval Hungarian elite from the Székesfehérvár royal necropolis

**DOI:** 10.64898/2026.04.10.717699

**Authors:** Oszkár Schütz, Zoltán Maróti, Kitti Maár, Luca Kis, Bence Kovács, Alexandra Ginguta, Balázs Tihanyi, Orsolya A. Váradi, Emil Nyerki, Petra Kiss, Monika Dosztig, Balázs Kertész, Frigyes Szücsi, Zoltán Szabó, Judit Olasz, Peter L. Nagy, Endre Neparáczki, Tibor Török, Gergely I. B. Varga

**Author notes:** These authors contributed equally to this work (Oszkár Schütz, Zoltán Maróti). City Archive and Research Institute, Székesfehérvár, Hungary (Balázs Kertész).

## Abstract

We analyzed 399 shotgun genomes from the Royal Basilica of Székesfehérvár, representing the elite of the Medieval Hungarian Kingdom. Our results reveal that the Carpathian Basin population underwent a marked homogenization during the Middle Ages, driven largely by admixture with eastern immigrant groups, especially the conquering Hungarians, resulting in a genomic landscape distinct from that of the surrounding European populations. The European ancestry component also shifted: the Avar-period Balkan element disappeared, while northern and northwestern European ancestry increased, likely reflecting changing political connections. We identified a previously unrecognized Conquest-period stratum at the site, including two individuals closely related to the Árpád dynasty. No large, continuous kinship networks were detected, indicating a dynamic and continuously renewed elite. At the population level, the strongest affinities were observed with the Conq_Asia_Core group and its ancestor, the Karayakupovo horizon, as well as with medieval populations from neighboring regions (present-day Croatia, Serbia, Slovakia, Poland, and Montenegro), reflecting the diverse sources of the medieval Hungarians.

## Introduction

The Church of the Virgin Mary in Székesfehérvár – commonly known as the Royal Basilica – was founded before 1031 by the first Hungarian king, Saint Stephen I (1000/1001–1038), as his private chapel. The chapter composed of canons developed from the clerical body attached to the church, headed by the provost. After the death of its founder, the basilica became the coronation church of the Kingdom of Hungary until the city fell to the Ottomans in 1543. The building served as the final resting place not only for several monarchs, their spouses, and numerous relatives but also for secular nobles, provosts, and presumably other members of the chapter (*1–4*).

The first known individual buried in the basilica was Prince Emeric, the son of King Stephen I, who died in 1031. Of the 37 Hungarian monarchs who died before 1543, a total of 15 are known to have been buried in Székesfehérvár. It is also evident that many other members of royal families were interred there, although the surviving records are incomplete (*1*, *4*). From at least the early 14^th^ century onward, both written records and archaeological evidence indicate that secular nobles and their families could also be buried in the Church of the Virgin Mary. Ecclesiastical figures who ended their lives as heads of the chapter were also buried here. Of these, only a fragment of the tombstone of Provost Miklós Györgyi Bodó (1444–1474) has been recovered (*1*, *5*). The last known person buried in the basilica was King John Szapolyai (d. 1540) but following the Ottoman conquest of the city in 1543, his body was removed from the church. Afterward, no further burials took place, and the basilica gradually fell into ruin.

Following the discovery of the burials of King Béla III (1172–1196) and his first wife, Anna of Antioch (d. around 1184), in December 1848, the basilica site was subject to multiple waves of excavation until the beginning of the 21^st^ century. These investigations uncovered the ruins of the building and revealed more than 900 unidentified human remains, two-thirds of which were found outside the structure (*6*, *7*) (Figure 1A). Several construction phases of the edifice have been identified, beginning with the earliest component, dating to at least the early 11th century, and concluding with a 15th-century late Gothic extension (*8–11*). Along with the royal couple, a small number of other excavated skeletons (indicated within our dataset with the ID “HU”) were transferred to Budapest, where they are now housed in the Matthias Church, while the majority of the human remains were deposited in the ossuary constructed at the basilica site, today known as the Medieval Ruin Garden, as well as in the depository of the Szent István Király Museum in Székesfehérvár. As the entire cemetery had been disturbed, there were few reliable archaeological indicators for dating the burials. Based on a limited number of finds with ambiguous chronology and stratigraphic observations, some graves have been proposed to date to the late 10^th^ century. However, these attributions have never been confirmed by additional evidence or radiocarbon dating, and several were later reassessed as 11^th^-century burials (*8*, *10*, *12*).

**Figure 1.**
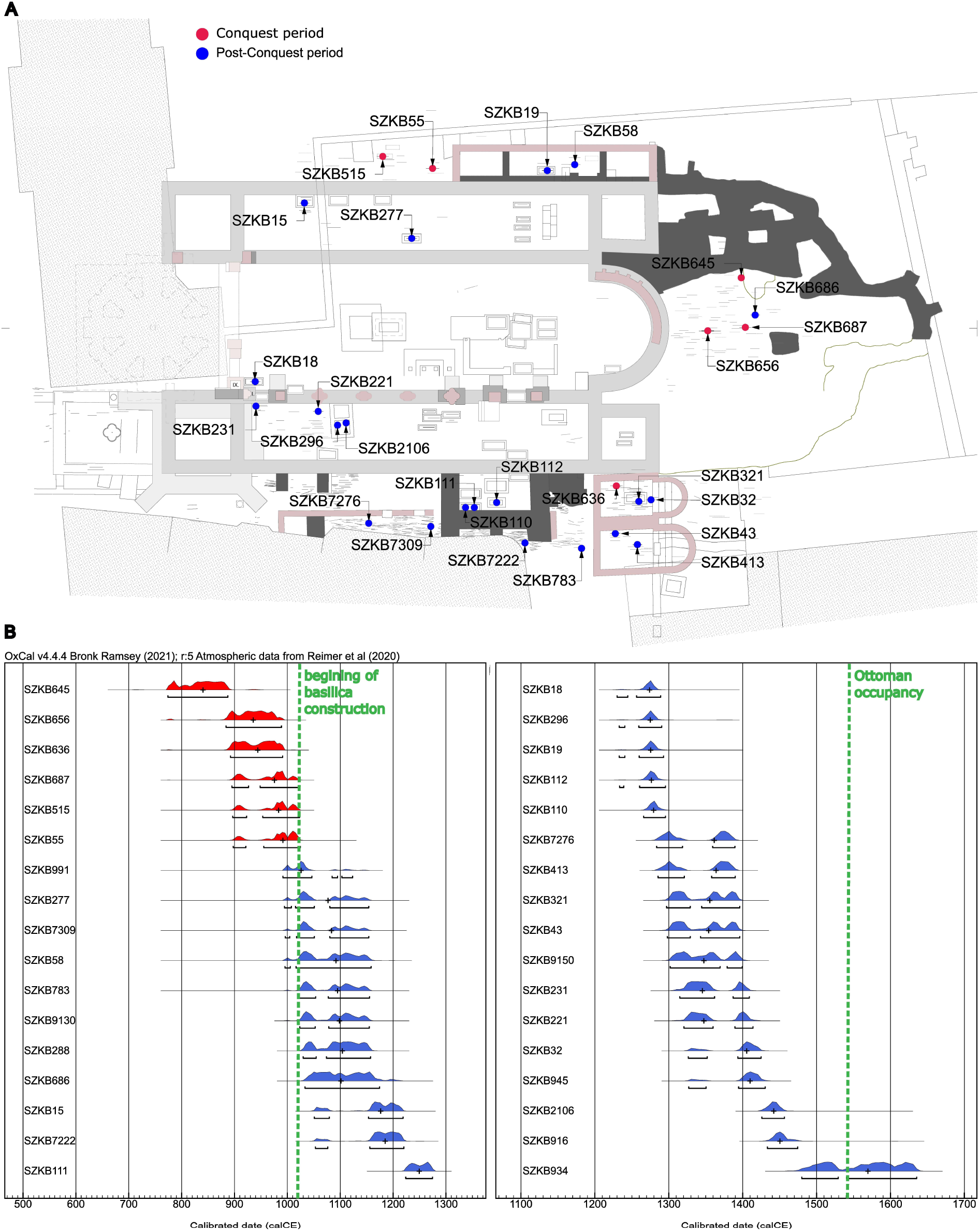
The reconstructed location of the radiocarbon dated remains at the Coronation Basilica site. **A**) The reconstructed ground plan of the Coronation Basilica and the location of the radiocarbon dated remains. Red dots indicate individuals buried before the basilica’s construction, while blue dots indicate those buried afterward. The burial locations of other human remains are indicated by thin gray lines. **B**) Age of the radiocarbon dated remains. Green dashed lines indicate significant boundaries in the history of the cemetery: the beginning of the basilica construction (1018) and the end of its use following the Ottoman occupation (1543).

A comprehensive genetic analysis of the remains associated with the coronation basilica offers a unique opportunity both to identify the royal individuals (*13*, *14*) and to characterize the genomic landscape of the medieval Hungarian elite between the 11th and 16th centuries. While the genomic history of earlier periods in the Carpathian Basin is relatively well understood, the events following the 10th-century Hungarian Conquest remain poorly explored. Population genetic studies have extensively investigated the early medieval Carpathian Basin. Several 6^th^ century Langobard and other Germanic sites from the Transdanubian region near Lake Balaton were analyzed, with probable origins traced to northern Europe (*15*, *16*). The geographic origins and core population groups of eastern steppe immigrants have also been examined in other studies. Using both statistical (*17*) and identity-by-descent (IBD) approaches (*18*), it has been shown that the military elite of the European Huns was genetically related to the Xiongnu of Mongolia, whereas the leaders of the Avar Khaganate likely derived from a Rouran-related gene pool prevalent in Mongolia and eastern Asia (*17*, *19*).

Genomic analyses of Hungarian Conquest-period individuals from the Carpathian Basin, together with Iron Age and medieval skeletons from the Ural region, reveal the genetically diverse and plausibly multi-ethnic origins of the conquering Hungarian tribes. A group of trans-Uralic genetic origin has been identified as a core population of the conquerors (*17*), which shows close genetic and genealogical connections to individuals of the Karayakupovo horizon in the 11–12th century southern Urals (*18*). Another stratum of the Conquest-period immigrants lacked Uralic ancestry and instead exhibited a Central–Inner Asian origin (*18*), from a source genetically similar to the Xiongnu (*17*). All studies consistently indicate that the basal population with a European genomic profile remained the majority in the Carpathian Basin throughout of the migrations, repeatedly overlaid by incoming eastern immigrant groups (*17*, *20*).

The study of the remains from the royal necropolis may help to address several key questions. What relationship existed between the immigrant elite of the 10th-century Hungarian Conquest period and the ruling strata of the medieval Hungarian Kingdom? To what extent did the Conquest-period steppe-related and the local European genetic components persist after the 10th century? Did the composition of the local European component change over time? Can the appearance of new genetic components be detected? Furthermore, how endogamous was the ruling elite? Is it possible to identify large, continuous kinship networks, or were such patterns absent due to the continuous renewal and replacement of the elite?

## Results

### Dataset and aDNA authentication

Our aim was to generate shotgun genome sequences from petrous bones or teeth from as many individuals as possible. Éry and colleagues sorted the remains of at least 935 individuals (*6*), but only 602 of these remains included skulls or skull fragments. We extracted DNA from all 602 remains, and from skeletal bones of 14 additional individuals (Table S1a). The dataset was further complemented by 9 additional individuals originally excavated at Székesfehérvár but later reinterned in the Matthias Church in Budapest. Six of these samples had previously been analysed and their Y-chromosome data reported (*21*), however we have completed their full genomic analysis here. Finally, 443 individuals with sufficient endogenous DNA content were sequenced using shotgun sequencing to an average genome coverage of 1.85x (range: 0.01x-16.64x).

Quality control with Schmutzi (*22*) and ANGSD (*23*) indicated negligible contamination in most samples (Table S1b). Forty-three individuals were excluded due to high contamination and one due to coverage below 0.25x, leaving 399 genomes for downstream analyses (Table S1b). For some of the analyses, we included previously published, identified Árpádian kings from Székesfehérvár, as well as individuals associated with the Árpád dynasty published in (*14*, *24*).

More than three-quarters of the individuals (76%) were genetically male (Table S1b), which is in good agreement with previous anthropological estimates (71%) (*6*). The predominance of male individuals is not unexpected, as the political and ecclesiastical leadership of the medieval ruling class was composed almost exclusively of males. This pattern is consistent with the interpretation that the basilica and its surroundings primarily functioned as a burial place for members of the elite.

### There is a Conquest-period stratum of the site

To characterize overall genetic variation in our medieval dataset, PCA was conducted with the Székesfehérvár genomes (SZKB) projected onto a background of present-day European individuals (*25*), (Figure 2A, Table S2a). Most samples form a continuous cluster that partially overlaps the PCA space of modern Central and Southern Europeans (including Hungarians, Croatians, and Romanians), yet the vast majority are shifted outside the range of present-day Europeans. The samples are also clearly distinct from previously published contemporary medieval European genomes (Figure 2B, Table S2b), including those from Poland (*26*), Scandinavia (*27–29*), Croatia (*30*), Italy (*27*, *31*, *32*), and Serbia (*30*). Exceptions include contemporaneous individuals from other regions of Hungary (*17*, *20*), two outliers from medieval Poland (*26*), and several medieval individuals from Montenegro (*32*). In addition, a significant proportion of SZKB genomes forms a clearly distinguishable genetic cline, shifted upward from the main dense cluster in Figure 2A.

**Figure 2.**
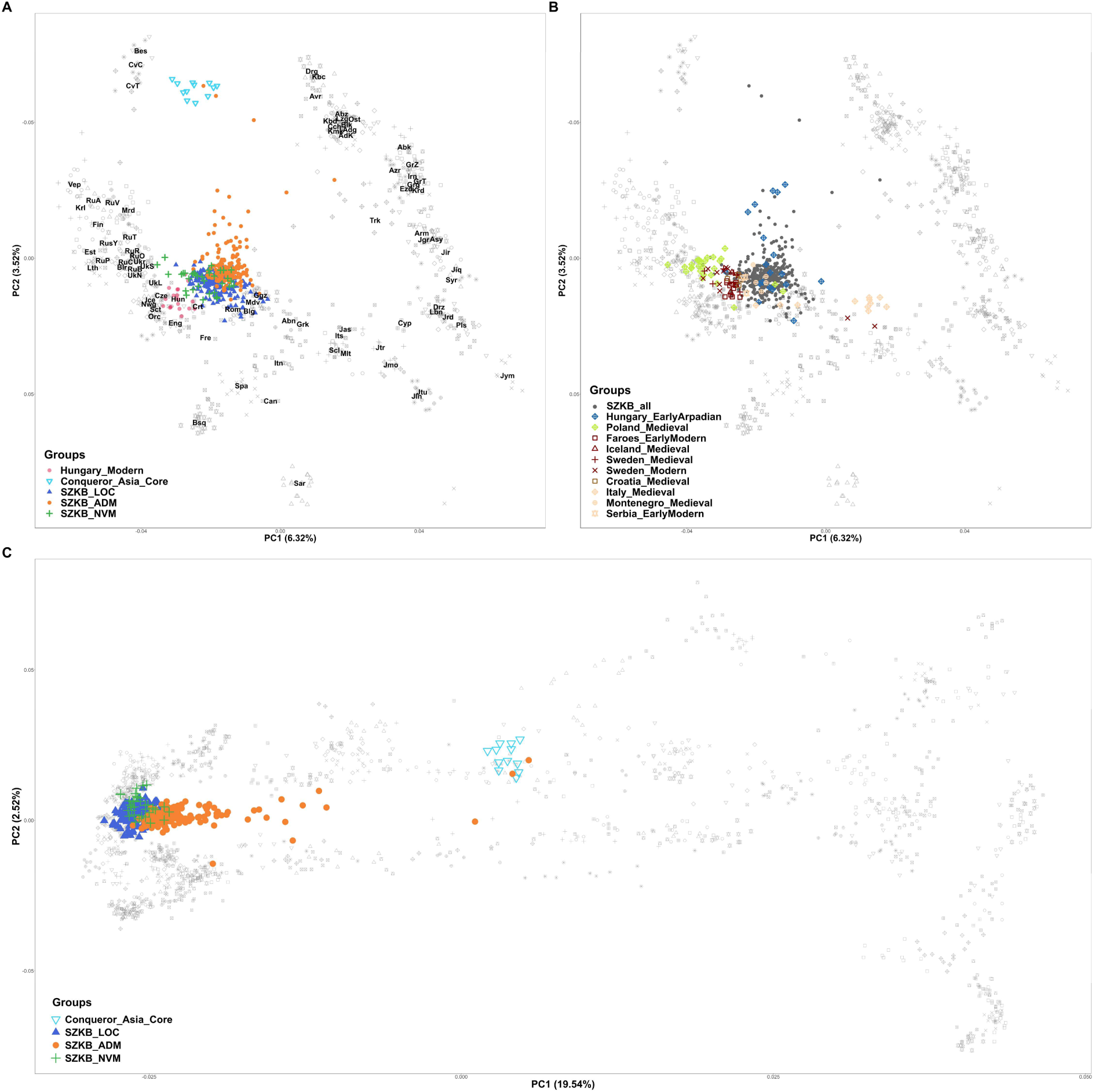
PCA of the remains excavated at the Coronation Basilica of Székesfehérvár. **2A:** SZKB genomes projected onto a reference panel of present-day European individuals (*25*), modern Hungarians are highlighted with red dots. The three-letter abbreviations of the populations are labeled at the median of their distributions. Conq_Asia_Core genomes from (*17*) are indicated by light blue inverted triangles. Additional colors and shapes denote groups based on qpAdm modeling: dark blue triangles - genomes modeled from local sources; orange circles - genomes with significant eastern components; green plus signs - genomes without a valid qpAdm model. **2B:** Published medieval genomes from contemporary European populations (Table S2b) projected onto the same European background as in Figure 2A. SZKB genomes are shown as dark gray dots for comparison. **2C:** SZKB genomes projected onto a background of present-day Eurasian populations (*17*).

When projecting the SZKB genomes onto a background of modern Eurasian populations (Figure 2C, Table S2c), this genetic cline is clearly shifted toward Asia and largely overlaps the “Conqueror cline” described in (*17*). The two easternmost individuals on PC1 (SZKB645 and SZKB687) fall squarely within the variance of the first-generation Conquering Hungarians, referred to as the Conq_Asia_Core in (*17*) suggesting that conquerors and their descendants compose the cline.

Radiocarbon dating of six samples indicates that these individuals were buried before the construction of the basilica (Figure 1B, Table S1a, Figure S1). All six samples fall into the “Conqueror cline” in PCA (Figure S2), with the earliest-dated individual, SZKB645, clustering with SZKB687 within the Conq_Asia_Core group (Figure 2A and 2C). As shown below, these later samples formed a clade with Conq_Asia_Core in qpAdm models and shared a substantial amount of IBD with all members of the Conq_Asia_Core group. These results provide clear evidence of a Hungarian Conquest-period layer in the cemetery. Since radiocarbon dating was performed on only a relatively small number of individuals, we cannot precisely distinguish all Conquest-period burials, however, most of them likely fall along the PCA cline. Notably, some of the radiocarbon-dated individuals buried after the basilica’s construction also fall within the cline (Figure S2).

The PCA distribution of the samples suggest that most individuals buried at Székesfehérvár represent admixture between the conquering Hungarians (or other groups with eastern genomic components) and the local population, with already diluted Asian genomic components. Compared to the 10th-11th century Conquest-period population (*17*), the genetic composition of the SZKB population appears much more homogeneous. This pattern likely reflects the homogenization of Asian and European ancestry components, explaining the clear separation of most SZKB genomes from both present-day and medieval European populations.

### Eastern ancestry largely derived from conquering Hungarians

To investigate the origin of the Asian genetic component, we screened each genome for the presence of significant Asian ancestry and evaluated whether it could be modeled from conquering Hungarians or other source populations using the qpAdm method (*33*). We tested two-source admixture models using European and Asian proxies, and applied the model competition approach (*17*) in order to narrow the list of valid models (Table S3). To model the local European ancestry components, we used the Eur_Core groups as proxies from Maróti (2022) (*17*), while the Asian components were represented by the Asia_Core groups of the known eastern immigrant populations; Hun_Asia_Core, Avar_Asia_Core, and Conq_Asia_Core from the same publication. We also aimed to model the eastern immigrant groups arriving after the Hungarian Conquest, such as the Cumans and Alans. As no genomes from these populations are currently available, we used the Russian_Alan group (*34*), as a proxy for the Alan source and the Ukraine_Cuman group together with the Kazakhstan_Kipchak1 and Kazakhstan_Kipchak2 genomes from the steppe published in (*34*, *35*) as proxies for Cuman-related ancestry.

The vast majority of genomes could be successfully modelled using these sources (Table S3), allowing us to divide the individuals into two groups: those yielding models exclusively from Eur_Core sources without significant eastern ancestry (SZKB_LOC, TableS3b, dark blue triangles in Figure 2), and those requiring eastern source in their models (SZKB_ADM, TableS3c, orange circles in Figure 2). The PCA positions of the individuals correspond well to their qpAdm models, although there is a narrow zone of overlap between the two groups. In this overlapping region, eastern ancestry may be too low for qpAdm to detect in all cases. Only a small number of genomes (n = 36) lacked a significant model, or had all significant models excluded during the model competition run (SZKB_NVM, Table S3d; green plus signs in Figure 2). The most likely explanation for this is the absence of their optimal source in the Left populations. Nevertheless, based on PCA, most of these genomes resemble the local population, likely with minor contributions from neighboring regions.

A total of 164 genomes showed significant eastern affinity, with most (n = 144) modeled from Conq_Asia_Core (Figure 3A). Many individuals could also be modeled with multiple eastern sources. The presence of these alternate models suggests that eastern ancestry is broadly shared across the applied immigrant proxy groups, which probably makes it difficult for qpAdm to discriminate sources, particularly when the ancestry proportion is low. Nevertheless, nearly two-thirds of the 164 individuals (n = 98) exhibit IBD connections with the Conq_Asia_Core group or its immediate ancestors, the Karayakupovo horizon (*18*) (Figure 3, Table S4). Based on this it is highly likely that the conquering Hungarians were the primary source of eastern ancestry in most cases. Notably, in some individuals a significant proportion of Conqueror ancestry persisted until at least the late 14th century. For example, individual SZKB221 (calibrated age 1320–1360 CE, 62.3%; 1389–1414 CE, 33.2%) carried 14% Conq_Asia_Core ancestry and exhibited direct IBD connections to the Conq_Asia_Core group.

**Figure 3.**
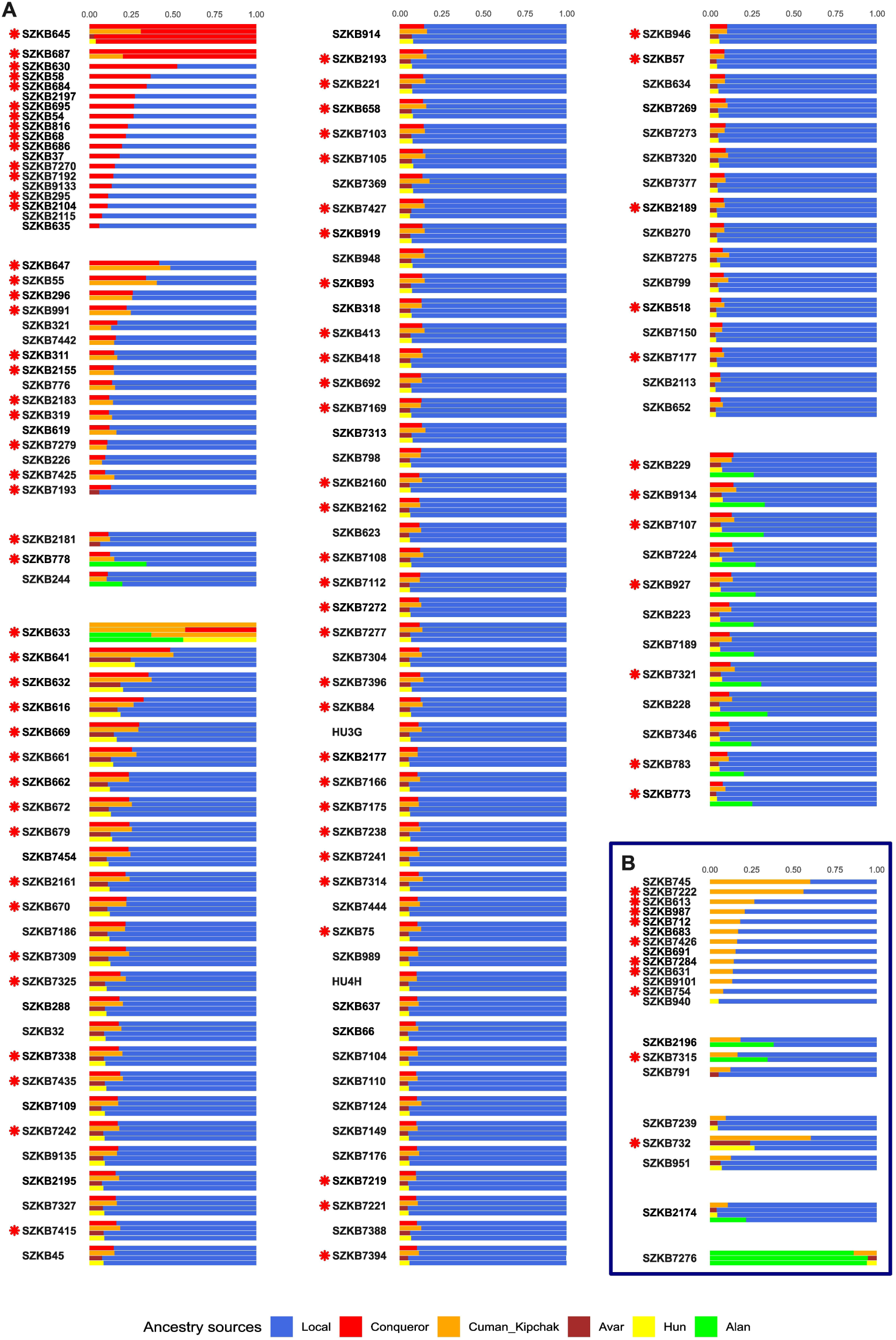
The qpAdm models of the genomes with eastern affinity. **A)** Genomes that could be modelled using the Conqueror Asia Core source. **B)** Genomes that could not be modelled using the Conqueror Asia Core source. The figure was created from the data in Table S3C. All alternative models are also shown using the sources indicated by the color codes at the bottom of the figure. Individuals sharing IBD with Conq_Asia_Core and/or Karayakupovo horizon (Table S4) are indicated by red asterix.

Four of the SZKB genomes did not bear a significant amount of local ancestry: two of these were the above-mentioned Conquest-period individuals, SZKB645 and SZKB687, who formed a clade with Conq_Asia_Core (Figure 3A). Moreover, they shared a significant amount of IBD with all members of the Conq_Asia_Core group (Table S4) and mapped among them on PCA. For 17 additional samples, all alternative models could be excluded, and the genomes could be modeled exclusively as a two-way mixture of Conq_Asia_Core and local populations. Importantly, 12 of the 17 samples also shared IBD with Conq_Asia_Core, corroborating the qpAdm results (Figure 3, Table S4).

Of the 164 samples with eastern affinity, only 21 produced models in which Conq_Asia_Core was rejected as a source during model competition, and in most of these cases, a Cuman/Kipchak source remained the Asian component (Figure 3B). Nonetheless, nearly half of these samples (n = 10) shared a significant length of IBD with Conq_Asia_Core, indicating that at least some of their eastern ancestry may in fact derive from the conquering Hungarians, even if qpAdm did not recover them as a source, likely due to the significant similarity of the Conq_Asia_Core and Cuman/Kipchak genome structures. Nevertheless, the high number of individuals that could be modeled solely as a mixture of local and Cuman-like ancestry (Figure 3B) suggests that descendants of later migration waves may also have been buried in the cemetery. A notable case is SZKB7276, whose ancestry could be modeled as 80–95% Alan-like, potentially representing the Jassic (Jász) people of Alan origin. This is consistent with the sample’s PCA position, as it is the only outlier projecting toward the Caucasus (Figure S2), in agreement with the reported PCA positions of the Alans (*34*); moreover, its radiocarbon date (Figure 1B) corresponds to the arrival of the Jász population (13th–14th centuries).

Based on these data, the most plausible conclusion is that conquering Hungarians were buried at the territory of the basilica before the church was founded. After the establishment of the site, members of the Hungarian Kingdom’s elite were buried here, most of whom were descendants of the conquering Hungarians already admixed with the local population. Some genomes with eastern components may also represent descendants of later migratory waves (i.e., Pecheneg, Cuman and Alan).

### Shift in local European ancestry

The qpAdm analysis identified 197 individuals lacking eastern ancestry (Table S3b), all modeled exclusively using Eur_Core proxies from Maróti et al. (2022) (*17*). The five Eur_Core groups from the 6th–11th centuries defined a north-south European genetic cline, highlighting the presence of nearly all major European genome types in the Carpathian Basin during this period. Using f4 statistics and hierarchical clustering, Gerber et al. (2024) (*20*) refined the Eur_Core classification of Maróti et al. (2022) (*17*) into four groups: a Near Eastern-Balkan-related CB-Eur group 1, a Northern European-related CB-Eur group 2, and two subgroups (3.1 and 3.2) representing a mixture or cline between them.

Our new dataset provided an opportunity to examine whether the composition of the local European ancestry changed during the 11th–16th centuries compared to earlier periods. Accordingly, all 197 European genomes and 36 additional samples lacking a significant model, or with all models excluded during model competition (Table S3d), were analyzed using the f4-statistics–based hierarchical clustering approach of Gerber et al. (2024) (*20*) (Table S5). First, we calculated the same three f4 statistics as in Gerber et al. (2024) (*20*) for all 197+36 genomes (Table S5a), and subsequently assigned the samples to the Gerber-defined clusters (Table S5b), based on the f4 values as described in the Methods.

All genomes could be assigned to one of the clusters, except for seven, which were considered as outliers. Based on the clustering results, we generated 2D plots using principal components derived from the f4 values, allowing a comparison of the distribution of SZKB genomes from the 11th–16th centuries with those from the earlier 6th–11th centuries. (Figure 4).

**Figure 4:**
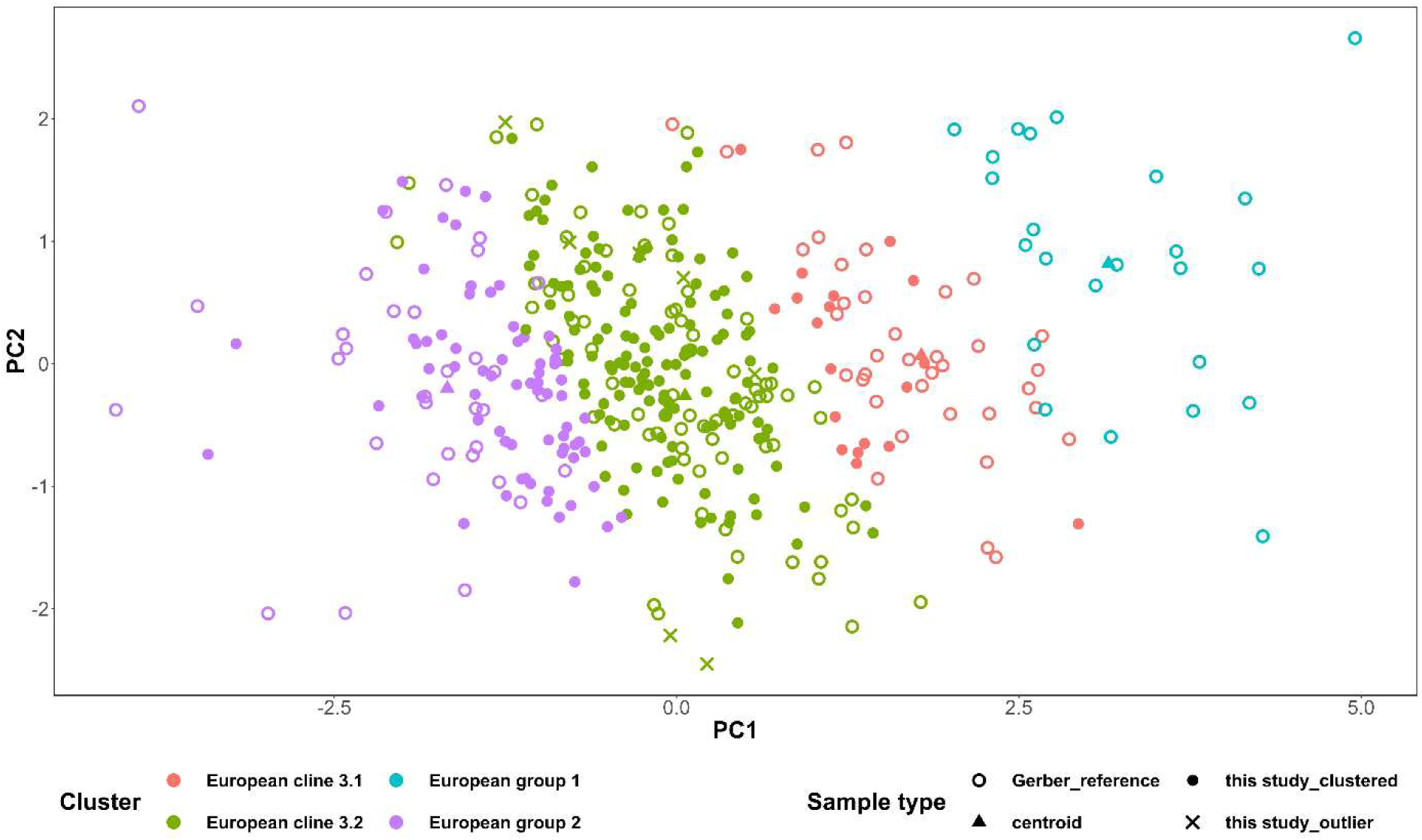
PCA of European genomes based on f4 values. Open circles indicate 6th–11th-century genomes from Gerber et al. (2024) (*20*), while filled circles represent 11th-16th century SZKB genomes. Blue denotes Group 1 (Balkan-associated), purple Group 2 (Northern European-like), red subgroup 3.1, and green subgroup 3.2. Triangles indicate the centroids of the four groups, and green crosses indicate the 7 outlier individuals.

The SZKB genomes form a markedly tighter cluster than their 6th–11th-century counterparts, with most individuals falling within the intermediate 3.2 cluster, indicating pronounced genetic homogenization in the Carpathian Basin during the Classical Medieval period. Notably, none of the SZKB genomes fall into CB-Eur Group 1, which predominantly reflects Near Eastern, Late Antique, and/or Balkan ancestry, particularly characteristic of the Avar period. This absence indicates a genetic shift toward affinities with Northern and Western Europe.

### Absence of large kinship networks in the Medieval elite

Internal genealogical connections were reconstructed using two complementary approaches: correctKin (*36*), which enables the identification of kinship relationships up to the 4th degree, even from low-coverage ancient genomes, and IBD analysis (*37*, *38*), which has a broader detection range extending beyond the 8th–9th degree, although precise estimation of the degree of relatedness is more challenging due to the stochastic nature of recombination. Given that the coverage threshold for correctKin is approximately 0.1x, 404 genomes could be included in this analysis (Table S6), whereas 395 genomes met the requirements for imputation and ancIBD analysis (Table S4). To assemble a comprehensive dataset from the site, previously published genomes from Varga et al. (2023) (*24*) and Kovács et al. (2026) (*14*) were also incorporated. The total shared IBD lengths and inferred kinship coefficients showed strong concordance in all cases, providing mutual validation of the inferred relationships. Both approaches also identified 7 duplicated individuals (Table S6), likely resulting from historical mix-ups of remains during excavation and storage (*6*, *7*). Among these, a highly plausible pair of monozygotic twins was also identified (SZKB321 and SZKB9135), as both DNA extracts were obtained from the left petrous bone, thereby excluding sample duplication.

Considering all pairwise IBD segments ≥8 cM, the SZKB genomes form a sparse population-level network (Figure S3). Among the 395 imputed genomes, 385 showed at least one internal IBD connection, yielding a total of 1,191 pairs; however, this represents only 1.37% of all theoretically possible internal connections. Moreover, 64% of the detected connections are limited to below 12 cM in length (77% below 16 cM), further underscoring the sparsity of the network.

Close genealogical relationships (1st–3rd degree) were rarely detected by either method. CorrectKin identified 38 such connections (TableS6), while ancIBD detected 33 pairs with total shared IBD >800 cM (Figure 5A, TableS4). More distant kinship relationships (4th–7th degree) were also uncommon: ancIBD identified 52 additional connections (TableS4) with shared IBD >50 cM, roughly corresponding to kinship up to the 7th degree. However, at such distances IBD sharing exhibits substantial variance, making it impossible to infer the exact degree of relatedness from a single IBD estimate.

**Figure 5.**
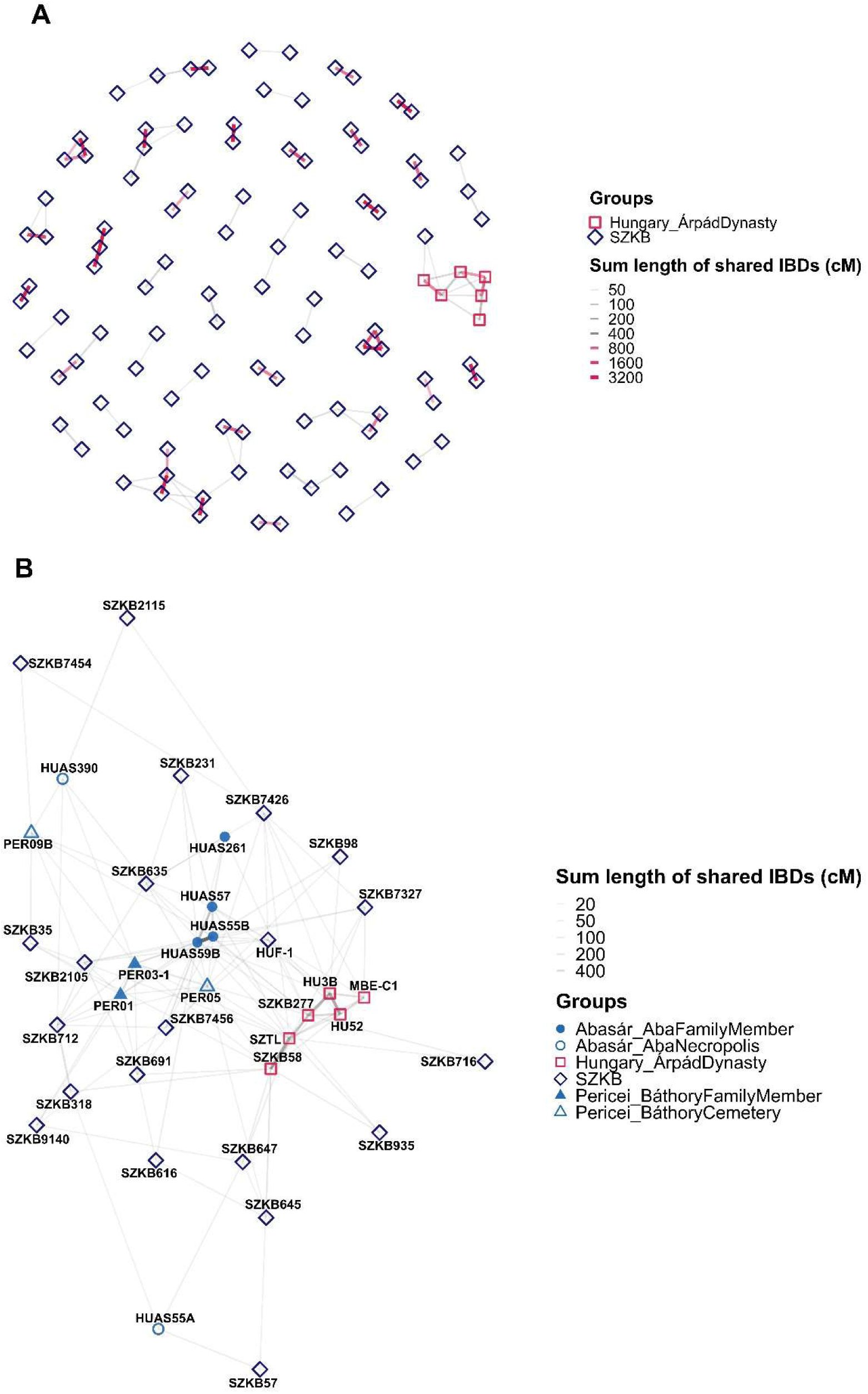
Internal IBD network of the necropolis. **A)** IBD network of close genealogical connections within the necropolis. For clarity, only shared IBD segments with a minimum total length of 50 cM are shown, with connections exceeding 800 cM highlighted in red. Previously identified members of the Árpád dynasty are also included. **B)** SZKB individuals with >20 cM IBD connections with the previously analyzed medieval Hungarian noble families, the Árpáds, Abas and Báthorys.

The most extensive kinship network, including both close and distant relatives, was observed among members of the Árpád dynasty and their relatives (Figure 5B) (*14*, *24*). We previously demonstrated that the early Árpád dynasty genomes (SZKB58 and SZTL) can be modelled as deriving from Conq_Asia_Core sources using qpAdm, and we also identified IBD connections linking them to this group (*14*, *24*). Our current results further reinforce this genealogical relationship. Notably, SZKB645, the earliest skeleton in the cemetery, and a representative of the Conq_Asia_Core group, shares 29 cM of total IBD with SZKB58, the earliest identified member of the Árpád dynasty (Figure 5B, Table S4). SZKB647, a female modelled as 42% Conq_Asia_Core, was excavated in close proximity to SZKB645 and thus likely belongs to the earliest burial horizon; she shares 76 cM with SZKB58 and 57 cM with SZTL. The SZKB935 male with unknown burial place shares 32 cM with SZKB58 and 25 cM with SZTL. Together, these findings provide direct genetic evidence of a genealogical link between the early Árpáds and Conqueror-related individuals buried at the site.

Several SZKB individuals show connections to later members of the Árpád dynasty as well as to other medieval noble families. SZKB716 (a male with local genome) shares 31 cM with SZKB277 and 8 cM with SZTL. SZKB7327 shares 23 cM with Béla III (HU3B) and 23 cM with Béla of Macsó (MBE-C1), and is also connected to the Aba (HUAS) family. Furthermore, 14 SZKB individuals exhibit IBD connections with previously analyzed medieval Hungarian noble lineages, including the Árpáds, Abas, and Báthorys (Figure 5B, Table S4).

### Group-based IBD sharing

To explore the broader population affinities of the SZKB group, we tested its pairwise group connectedness with 64 other medieval European and relevant Asian populations, based on a compiled dataset of published genomes dating from the 5th century onward (Table S7). For this analysis, we divided the SZKB population into three subgroups based on their qpAdm-inferred genomic composition: (1) individuals with exclusively European ancestry (SZKB_LOC), (2) individuals with detectable Asian ancestry components (SZKB_ADM), and (3) individuals for whom no valid qpAdm model could be obtained (SZKB_NVM).

To evaluate connectivity between group pairs and enable comparability across groups we normalized the group degree centrality by the total number of possible connections, resulting in the ratio of fulfilled connections (*38*). We calculated this index-number for both inter-group and intra-group connections of the SZKB population and its subgroups (Table S7).

The external IBD connections of the SZKB population (Figure 6 and Table S7) are in good agreement with the qpAdm results. The SZKB_ADM subgroup, which carries Asian ancestry components, exhibits an intra-group fulfilled connection value of 0.022, which is nearly twice as high as that of the SZKB_LOC (the subgroup modelled exclusively from European genomes 0.0137) or the combined SZKB group (0.0137) (Table S7). This suggests that SZKB_ADM forms a tighter cluster, likely because of a shared Asian ancestry component derived from a common source. As the strongest connections of the SZKB_ADM subgroup were observed with the Conq_Asia_Core (inter-group fulfilled-connection value; 0.092) and the Karayakupovo horizon (0.048), this strongly indicates that the shared source is most likely linked to the conquering Hungarian population.

**Figure 6.**
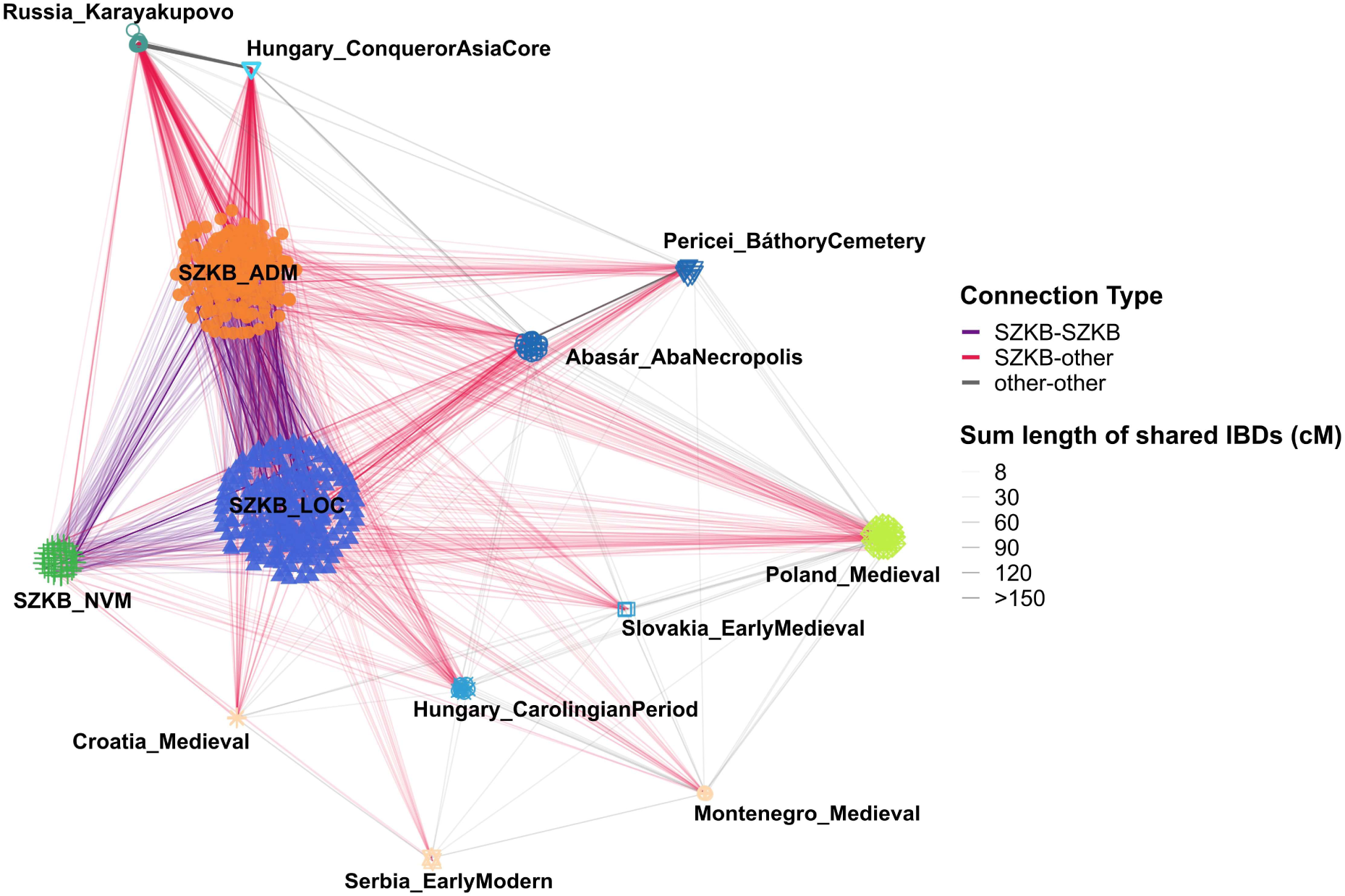
Major external population connections of the SZKB subgroups. The SZKB population was divided into three subgroups, as detailed in the text. Only those groups are shown in the figure that exhibit a higher connection rate with at least one of the SZKB subgroups than the within-group connection rate of the given SZKB subgroup (Figure S4). For the complete group-based data, see Table S7.

The SZKB_ADM subgroup also shows significant relatedness to several medieval groups from Hungary, including Caroling era population (*17*, *20*), members of the Báthory (*25*) and Aba (*39*) families, as well as groups from the broader Carpathian Basin region, such as Slovakia_EarlyMedieval and Croatia_Medieval (*30*).

The strongest connections of the SZKB_LOC subgroup closely mirror those observed for SZKB_ADM (Table S7b and S7c): the highest fulfilled-connection values were detected with the same Carpathian Basin and surrounding Central European groups listed above, as well as with Roman-period samples from Montenegro, and Ottoman-period samples from Serbia (*32*), and, notably, with Conq_Asia_Core as well (Figure 6, Table S7, Figure S5).

The SZKB subgroup lacking valid qpAdm models (SZKB_NVM) shows nearly the same external connections as the SZKB_LOC subgroup, with the exception of a weaker connection to Conq_Asia_Core and some exclusively higher connectedness to Medieval samples from Poland and Croatia (*26*). Notably, all SZKB subgroups exhibit very strong connections to one another, indicating that they effectively constitute a single, cohesive population. This interpretation is further supported by close familial relationships, including several cases of close kinship between individuals assigned to different subgroups (Figure S6).

## Discussion

In this study, we examine the genetic characteristics of the classical-medieval Hungarian elite (11th–16th centuries) through a comprehensive genome-wide analysis of a substantial and representative sample set. Although the 399 genomes analyzed likely pertain to the elite, it is plausible that the broader population of the Carpathian Basin during this period exhibited a similar genetic profile. Our findings indicate that the medieval Hungarian population differed markedly from most contemporary and present-day European populations, as evidenced by its distinct positioning in the PCA space. While it exhibits the closest genetic affinity to modern Hungarians and neighboring Central European groups including contemporaneous populations from present-day Croatia, and Romania, their detectably higher proportion of Asian ancestry distinctly sets them apart from other European populations.

While the Asian genetic component in the Carpathian Basin may have been introduced through multiple eastern migratory waves, such as the Huns, Avars, conquering Hungarians, Pechenegs, and Cumans, our data indicate that the genetic legacy of the conquering Hungarians was the most influential among these groups. Furthermore, our findings suggest that by the medieval period, the population of the Carpathian Basin had become significantly more homogenized compared to the 10th century. Consequently, the Asian ancestry component became widespread across a substantial portion of the population, albeit in a diluted form. Previously studied high-status individuals, including members of the Árpád (*14*, *24*), Aba (*39*), and Báthory (*25*) families, exhibit the same genomic structure. Our expanded dataset shows that this composition was not unique but rather widespread across the medieval Carpathian Basin, as further evidenced by contemporary Árpádian-period genomes excavated elsewhere, which fall into the same PCA distribution (Figure 2B). This homogenization process likely persisted in subsequent centuries, which may account for the observation that contemporary Hungarians still display a somewhat elevated proportion of eastern ancestry compared to neighboring populations (*40*).

A particularly novel finding is that radiocarbon dating and ancient genomic data consistently reveal a Conquest-period stratum in the cemetery that predates the construction of the basilica itself. Even more striking is that several individuals, including the earliest skeleton in the cemetery (SZKB645), exhibit direct genealogical links to the early Árpád dynasty. If the radiocarbon date of 774 (95.4%) 887 calCE for this skeleton is accepted, it suggests the presence of Hungarians in Eastern Transdanubia before the traditionally accepted date of 895 and 900, in line with recently published cases from Zalavár (*20*). This may suggest that the site already functioned as a burial place for relatives of the Árpád lineage from the time of the Hungarian Conquest, thereby shedding new light on the role of Székesfehérvár as a 10th-century power center of the Hungarian Principality. It remains an open question whether these Conquest-period burials were associated with an early church, possibly a predecessor of the later basilica (*8*, *10*, *41*).

The data further suggest that the genetic composition of the Carpathian Basin underwent dynamic changes throughout the medieval period. In addition to weak and uncertain signals of subsequent eastern migration waves (e.g., Pechenegs, Cumans, Alans), a discernible shift is evident in the European ancestry components. While earlier periods were characterized by a substantial Near Eastern–Balkan-related component - likely associated with the legacy of the Roman Empire - this component had largely disappeared by the medieval period, probably due to assimilation or substantial migration. Alternatively, it is possible that this southern-derived ancestry was underrepresented among the elite. In contrast, the proportion of northern European ancestry, which has long been characteristic of the region, remained relatively stable. These findings align with historical evidence indicating intensified connections with the Holy Roman Empire and other Western European polities following the foundation of the Hungarian Kingdom, including dynastic marriages, missionary activity, and the settlement of soldiers (*41–44*).

The relatively sparse internal IBD network of the SZKB genomes suggests that, while these individuals can be considered part of a broader metapopulation, close kinship ties were rare. This implies that those buried in and around the royal basilica constituted a socially constructed community rather than a tightly knit biological kin group. Taking into account the marked male bias, this pattern aligns well with expectations for a dynamically changing power structure. Our data suggest that the Hungarian elite of the 11th–16th centuries was not composed of long-standing dominant lineages, but rather was continuously reshaped, with new families regularly entering its ranks. This pattern may be particularly characteristic of those buried at Székesfehérvár; however, it is also possible that members of major noble families were interred in separate burial sites such as those of the Abas and Báthorys (*25*, *39*).

Finally, the external IBD network of the SZKB population closely reflects the inferred sources of the medieval Hungarian gene pool. The conquering Hungarians appear to have played a central role, through extensive admixture with the local populations of the Carpathian Basin. These strong IBD connections extend even to the Karayakupovo populations, representing the Uralic predecessors of the conquerors (*18*). At the same time, clear signals of admixture with surrounding populations are also evident. Most outlier individuals likely carry minor genetic components originating from neighbouring groups that were not represented in the earlier local European (Eur_Core) populations. The connections of all three SZKB subgroups with neighboring populations and medieval groups from Hungary are in line with expectations; however, the substantial IBD sharing between the SZKB_LOC subgroup and Conq_Asia_Core warrants explanation. The most parsimonious interpretation is that the ∼20% of SZKB_LOC individuals exhibiting IBD sharing with Conq_Asia_Core also harbor an Asian ancestry component, albeit at low levels. In reality, these individuals should have been assigned to the SZKB_ADM subgroup, however, we applied an arbitrary threshold and classified as SZKB_LOC all individuals for whom alternative qpAdm models yielded plausible fits based exclusively on Eur_Core sources. Accordingly, these individuals are expected to fall within the overlap between the SZKB_LOC and SZKB_ADM subgroups in PCA, which is indeed observed (Figure S5).

Our conclusions are constrained by the limited availability of radiocarbon data. Ideally, dating all samples would allow for a more precise delineation of the earliest Conquest-period section of the cemetery. Moreover, additional contemporary elite and commoner genome data from the Carpathian Basin and surrounding regions could further clarify the accuracy and broader applicability of our findings.

## Materials and Methods

### Sample selection

The Church of the Virgin Mary in Székesfehérvár was excavated on multiple occasions, and the recovered remains were extensively documented by Éry et al. (2008) (*6*) through detailed drawings, photographs, and comprehensive anatomical, anthropological, and pathological analyses. From the 935 individuals sorted by Éry and colleagues, we sampled the petrous bone or teeth of 602 individuals with preserved skulls or cranial fragments (Table S1a); the remaining individuals lacked cranial material.

In numbering the samples, we retained the original annotation system, in which the first digit indicates the burial context: 1 - interior stone grave; 2 - interior earth grave; 3 - Chapel 1; 4 - Chapel 2; 5 - northern exterior; 6 - eastern exterior; 7 - southern exterior; 8 - western exterior; 9 - unknown. The subsequent digits denote serial numbers. The original locations of the remains are shown in Figure 1A using thin lines.

In addition to the sorted remains of 935 individuals, the Székesfehérvár basilica assemblage also includes a substantial amount of unsorted, fragmentary mixed bone material, which was not included in our analyses.

We also included four additional remains excavated in Székesfehérvár but currently curated in the Matthias Church in Budapest (IDs: HU2per56, HU2per58, HU2per59 and HU6per1).

### Radiocarbon dating

Radiocarbon analysis was performed on the sampled bone fragments to confirm the archaeological dating of the remains. The measurements were done by accelerator mass spectrometry (AMS) in the AMS laboratory of the Institute for Nuclear Research, Hungarian Academy of Sciences, Debrecen, Hungary (AMS Lab ID: DeA-37107; technical details concerning the sample preparation and measurement: Molnár et al. 2013) (*45*). The conventional radiocarbon date was calibrated with the OxCal 4.4.4 software (https://c14.arch.ox.ac.uk/oxcal/OxCal.html, date of calibration: 9th of January 2024) with IntCal 20 settings (*46–49*).

### DNA extraction, library preparation and sequencing

All steps of sampling, DNA extraction, and library preparation were performed as described in the Supplementary Materials of Varga et al. (2023) (*24*), in the joint, dedicated ancient DNA facility of the Department of Archaeogenetics (Institute of Hungarian Research) and the Department of Genetics, University of Szeged. Double-stranded libraries were first shallow shotgun sequenced on an Illumina iSeq platform to assess human DNA content. Selected libraries above 10% endogenous DNA content were subsequently sequenced to greater depth on an Illumina NovaSeq platform, achieving an average genome coverage exceeding 1× for 443 samples (Table S1b).

An additional 100 samples with low human DNA content were enriched using the myBaits Expert Human Affinities Kit - Prime Plus (cat. no. 352096; Arbor Biosciences). These enriched genomes were excluded from the analyses due to their low quality. Genetic sex and haplogroup data for these samples are provided in Table S1c.

### Raw data handling, quality checking

The adapters of paired-end reads were trimmed with the Cutadapt software (*50*), and sequences shorter than 25 nucleotides were removed. Read quality was assessed with FastQC (*51*). The raw reads were aligned to GRCh37 (hs37d5) reference genome using the Burrows-Wheeler Aligner (v 0.7.17) software, with the MEM command in paired mode, with default parameters and disabled reseeding (*52*). Only properly paired primary alignments with ≥ 90% identity to reference were considered in all downstream analyses to remove exogenous DNA. Samtools v1.1 was used for merging the sequences for different lanes, sorting, and indexing binary alignment map (BAM) files (*53*). PCR duplicates were marked using Picard Tools MarkDuplicates v 2.21.3 (*54*). To randomly exclude overlapping portions of paired-end reads and to mitigate sequencing errors and potential random pseudo-haploidization bias, we applied the mergeReads task with the options “updateQuality mergingMethod=keepRandomRead” from the ATLAS package (*55*). Single nucleotide polymorphisms (SNPs) were called using the ANGSD software package (version: 0.931-10-g09a0fc5) (*23*) with the “-doHaploCall 1 - doCounts 1” options and restricting the genotyping with the “-sites” option to the genomic positions of the 1240K panel.

Ancient DNA damage patterns were assessed using the ATLAS package PMD task (*55*). Mitochondrial genome contamination was estimated using the Schmutzi algorithm (*22*). Contamination for the male samples was assessed by the ANGSD X chromosome contamination estimation method (*56*) as well, with the “-r X:5000000-154900000 -doCounts 1 -iCounts 1 -minMapQ 30 -minQ 20 -setMinDepth 2” options (Table S1b). We excluded samples above 4% contamination levels or below 0.25x coverage.

Raw genome data of six individuals from Nagy et al (2021) (*21*) were also included in the analyses (Table S1).

### Genetic sex and haplogroup determination, kinship

Biological sex was assessed with the method described in (*57*). Fragment length of paired-end data and average genome coverages (all, X, Y, mitochondrial) was assessed by the ATLAS software package (*55*) using the BAMDiagnostics task. Detailed coverage distribution of autosomal, X, Y, mitochondrial chromosomes was calculated by the mosdepth software (*58*) (Table S1b).

Mitochondrial haplogroup (Mt Hg) determination was performed with the HaploGrep 2 (version 2.1.25) software (*59*), using the consensus endogen fasta files resulting from the Schmutzi Bayesian algorithm (Table S1b). The Y Hg assessment was performed with the Yleaf software tool (*60*), updated with the ISOGG2020 Y tree dataset (Table S1b) (*61*).

Kinship analysis was performed with correctKin (*36*) (Table S6) on samples that had greater than 0.05 genotyping fraction, less than 5% contamination (according to mtDNA contamination by Schmutzi). All duplicate samples were excluded from the analysis. To ensure a reference population fitting well to our dataset according to PCA, qpAdm, IBD and chronological data, we applied the medieval genomes from Hungary published in (*14*, *17*, *24*, *25*, *39*, *62*). The meaning of the columns in Table S6 are as follows. **Uncorrected kinship coefficient:** raw pairwise kinship coefficient estimated between two samples, this value may be systematically biased downward in low-coverage datasets; **Calculated overlap fraction:** the marker overlap fraction between the two measured samples calculated by dividing the number of markers where both samples are genotyped with the total number of markers in the dataset (1240K); **Corrected kinship coefficient:** the estimated kinship coefficient divided with the marker overlap fraction; **6.00 sigma threshold:** a statistical significance threshold corresponding to six standard deviations from the null distribution of kinship values under the assumption of unrelated individuals. Values exceeding this threshold represent extremely strong evidence against the null hypothesis of no relatedness; **95 confidence interval (lower boundary):** the lower bound of the 95% confidence interval for the corrected kinship coefficient, representing the minimum statistically supported value of relatedness consistent with the observed data; **96 confidence interval (upper boundary):** the upper bound of the 95% confidence interval for the corrected kinship coefficient, representing the maximum statistically supported value of relatedness consistent with the observed data; **Estimated relatedness:** a categorical assignment of biological inferred from the corrected kinship coefficient using predefined threshold ranges.

### Principal Component Analysis

To uncover population structure based on genomic data, we conducted principal component analysis (PCA) (*63*). The Western Eurasian PCA was conducted as described in Patterson (2006) (*25*). For the background 784 suitable individuals belonging to 85 populations (primarily west of the Urals, with additional North African groups) were selected from the contemporary individuals present in the AADR dataset.

The Eurasian PCA followed Maróti (2022) (*17*), and included 1,397 modern individuals from 179 populations from the AADR data set that has PASS-ed quality assessments. To balance the dataset, generally no more than 10 individuals were selected per population including the samples with the highest genotyping rates.

PCA Eigen vectors were calculated from the 784 and 1397 pseudo-haploidized modern genomes, confined to the HO SNP set, with smartpca (EIGENSOFT version 7.2.1) (*63*). All ancient genomes were projected on the modern background with the ‘‘lsqproject: YES and inbreed: YES’’ options. Since the ancient samples were projected, we used a more relaxed genotyping threshold (>50k genotyped markers) to exclude samples only where the results could be questionable due to low coverage.

### qpAdm

We used qpAdm (*33*) from the ADMIXTOOLS software package (*64*) primarily for detecting significant Asian affinity in the studied genomes, and secondly for modelling our genomes as admixtures of two source populations and estimating ancestry proportions. The qpAdm analysis was performed using the 1240K dataset.

The Left population list was compiled to include available source populations representing both local European groups and medieval immigrants with Asian ancestry. For local European sources, we selected the Eur_Core1–5 groups from Maróti et al. (2022) (*17*), each reflecting slightly different genomic profiles from the 5th–11th-century Carpathian Basin. To capture eastern ancestry, we included Hun, Avar, and conquering Hungarian groups from Maróti et al. (2022) (*17*), Ukrainian Cumans from Saag et al. (2025) (*35*), and Kipchaks and Alans from Damgaard et al. (2018) (*34*). This selection aimed to encompass known sources with Eurasian Steppe and Caucasian ancestry associated with the medieval Carpathian Basin.

The reference population set (Right populations) was composed based on our previous experience in modelling steppe ancestry in the mixing population of the Hungarian Conquering Era (*17*). Thus, it included Ethiopia_4500BP (fixed), Iran_GanjDareh_N.AG, Romania_Mesolithic_IronGates.SG (WHG), Turkey_Marmara_N, Latvia_Mesolithic, EEHG, Tajik, Han, Mansi.DG. For the detailed list of Left and Right populations and the exact genomes included see Table S3.

During ‘base model strategy’ run we set the details:YES parameter to evaluate Z-scores for the goodness of fit of the model (estimated with a Block Jackknife). As qpWave is integrated into qpAdm, the nested p-values in the log files indicate the optimal rank of the model. This means that if the p-value for the nested model is above 0.05, the Rank-1 model should be considered (*33*).

In order to further exclude suboptimal models, we applied the ‘‘model-competition’’ approach described in Maróti et al. (2022) (*17*). For each individual tested, we have obtained a subset of the Left populations as possible sources present among the valid models. We moved the putative sources one at a time to the Right set and rerun qpAdm for each model. As the best sources have the highest shared drift with the Test population, including these in the Right list is expected to exclude all models with similar suboptimal sources. This way we were able to filter out most suboptimal models and identify the most plausible ones, which were not excluded by any of the Right combinations. In each run we considered only models with a p-value of at least 0.05 as valid. A valid model was excluded if any combination of the model-competition Right populations reduced its p-value below 0.05. We have also considered further optimality measurements indicated in the qpAdm output log. A “bad fitting” model was considered if the obtained source coefficients showed high standard error (>10%) and a “negative model” was considered if the source coefficients slipped out of the (0–1) conventional bounds.

The meaning of the columns in Table S3 are as follows. **Test:** the name of the individual modelled. **SourceX:** the name of the designated sources in descending order of contribution. **SourceX ratio**: the obtained mixture coefficient for the source population/individual in question. **Valid models**: the number of cases a given model passed the quality criteria. **Excluded models**: the number of cases a model has been excluded by one of the reference populations. **BadFit models**: quality criterion, number of cases the obtained mixture coefficient differs from the jackknife calculated mixture coefficient by at least 10%. **Negative models**: quality criterion, number of cases where one of the source coefficients is lower than 0. **Non-significant nested p-value models**: the number of models where the obtained nested p-value is higher than 0.05. This indicates that the k-1 model is more plausible. **Average nested p-value**: the average of the obtained p-values for the k-1 model. **Minimum p-value**: the lowest p-value obtained from the model competitions. **Maximum p-value**: the highest p-value obtained from the model competitions. **Average p-value**: the average of the obtained p-values of the valid model repetitions (passing the quality criteria). **Average p-value summary**: either the average p-value or the average nested p-value if the number of non-significant nested p-value models is higher than the half of the valid models. This metric was only used to order the qpAdm results. **Minimum p-value reference**: the name of the LEFT population/individual which was moved to the RIGHT population set when the lowest p-value was obtained for the model in question. **Maximum p-value reference**: the name of the LEFT population/individual which was moved to the RIGHT population set when the highest p-value was obtained for the model in question. **Excluding reference**: the name of the LEFT population/individual which was moved to the RIGHT population set when the model was excluded.

We applied the results of our qpAdm approach in two steps. First, we took advantage of qpAdm’s ability to measure genetic affinities simultaneously along several f4 axes and to integrate them into a single model. In this way, we distinguished between “local-type” and “admixed-type” genomes. A genome was classified as “admixed-type” (SZKB_ADM) if all valid models included an eastern source and no Rank-1 model based on a local (Eur_Core) source had a p-value > 0.05. Conversely, a genome was classified as “local-type” (SZKB_LOC) if it yielded at least one valid Rank-1 model based on a local source, or an admixture model consisting of two local sources.

Figure 3 was generated from Table S3c. All the genomes classified into SZKB_ADM were included. The sources were contracted into the following 6 groups: Local (Eur_Core 1-5), Conqueror (Conq_Asia_Core 1 and 2), Avar (Avar_Asia_Core1 and 2), Hun (Huna_Asia_Core), Cuman (Ukraine_Cuman, Kazakhstan_Kipchak1 and 2) and Alan (Russian_Alan). All the valid models of each genome after model-competition run were grouped according to the eastern source groups. The average of the ancestry ratios in different model types was taken. Finally, the average values were represented on the charts. The genomes were grouped on the figure according to their number of different model types. The figure was generated in R with the package *ggplot2* (*65*).

### Cluster assignment based on f4 statistics

To assign SZKB_LOC and SZKB_NVM individuals, likely lacking Asian affinity based on PCA and qpAdm, to the clusters published in Gerber et al. (2024) (*20*), we first replicated the clustering process of Gerber et al. (2024) (*20*) using the original dataset. For each cluster, we determined the centroid by calculating the median across all f4 dimensions to minimize the impact of potential outliers.

Next, we computed the same f4 statistics for each SZKB sample and calculated their Euclidean distance to each centroid 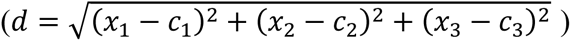 assigning each sample to the nearest Gerber cluster. This method successfully assigned all SZKB_LOC and SZKB_NVM genomes to a cluster. To detect potential outliers, we applied a threshold based on the 95th percentile of distances from the centroids, which resulted in the detection of seven outliers.

To visualize our results in Figure 4, PCA was performed on the f4-value matrix using the *prcomp* function (*66*) with the option scale. = TRUE. All individuals from the Gerber clusters and the European cloud were included. Figure 4 displays PC1 and PC2. Calculations and figure preparation were carried out in R (version 4.1.2) using the dplyr (*67*), tidyr (*68*), ggplot2 (*65*), and plotly (*69*) packages.

### IBD analyses

For imputation, we used the GLIMPSE2 framework (version 2.0.0) (*70*) using the 1KG Phase 3 dataset common markers as reference. The reference data set was normalized and multi allelic sites were split using bcftools (version 1.16-63-gc021478 using htslib 1.16-24-ge88e343) with the “norm –m –any” subcommand and filtered for biallelic SNPs with the “view –m 2 –M 2 –v snps” subcommand. The autosomal chromosomes of the human reference genome were divided into 580 genomic chunks using the GLIMPSE2_chunk tool with the “-sequential” option. As described in the GLIMPSE2 manuscript, we created the binary reference data with the GLIMPSE2_split_reference tool using the 580 genomic regions and the 1KG biallelic SNP variants.

In all downstream IBD analysis, we used only samples with >0.25x mean genome coverage of shotgun WGS data as recommended in the ancIBD manuscript (*37*). We excluded all samples with estimated MT contamination higher than 0.03 (based on the Schmutzi MT contamination analysis) (*22*) since the concordance of higher MT contamination (0.06-0.12) samples had significantly lower concordance according to our experiments using high coverage aDNA data (data not shown). Furthermore, we calculated the fraction of >0.99 GP (the variant genotype probability based on GLIMPSE2) fraction of variants of the imputed AADR chromosome 3 markers. Based on this value, we removed samples with less than 0.8 fraction of variants that had higher than 0.99 GP to exclude samples with excessive low confident data.

We used the ancIBD (version 0.7) python libraries with the Python 3.6.8 environment for IBD fragment analysis (*37*). Phased and imputed variants of experimental aDNA samples were post-filtered to include only the positions of the 1240K AADR marker set and lifted to the hdf5 data format as described in the ancIBD manuscript. IBD fragments were identified with the default parameters recommended for aDNA analysis (emission model haploid_gl2, HMM model FiveStateScaled, and the p_col=’variants/RAF’ option to use GLIMPSE2 reference AF data from the imputed variants). We identified IBD fragments >=8cM. During the subsequent filtering of raw IBD segments, we deviated from the marker density threshold (≥ 220 SNPs/cM) used in the original ancIBD framework and instead applied the marker informativity score approach described in Schütz (2025) (*38*, *71*), which provides greater power to detect true IBD segments.

Shared IBD networks were generated in R, with the application of packages *ggplot2* (https://ggplot2.tidyverse.org/) (*65*), and igraph (https://igraph.org/) (*72*, *73*), with Fruchterman-Reingold weight directed algorithm.

Group-based evaluation of IBD connections was conducted as described in Schütz et al. (2025) (*38*). To assess connectivity between group pairs and ensure comparability across groups, we normalized the total number of observed connections.

Let us define a subgraph of the IBD network in which vertices are selected based on a specified grouping parameter (here, the Fig. 6 label). Edges were then subsampled depending on the type of connectivity considered: for intergroup connectivity, only edges between vertices belonging to two different grouping categories were retained, whereas for intragroup connectivity, only edges between vertices within the same category were included.

The size of each subgraph, denoted as 𝑠_ij_, represents the number of connections between groups 𝑖and 𝑗. This value can be calculated as half the sum of vertex degrees within the subgraph:

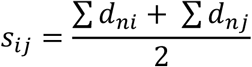

This measure serves as a natural descriptor of the connection strength between two groups. However, the maximum possible number of connections depends on the sample sizes (𝑛) of the groups:

- for intragroup connections: 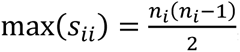,
- for intergroup connections: 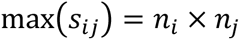.

Because these maxima increase exponentially with group size, direct comparison of raw connection counts is not intuitive. To address this, we normalized the observed number of connections (𝑠_ij_) by the corresponding theoretical maximum (max (𝑠_ij_)), yielding a normalized measure 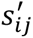 ranging between 0 and 1. This value represents the rate of fulfilled connections relative to all possible connections between the groups.

Figure 6 includes only those groups that exhibit a higher fulfilled connection rate with at least one SZKB subgroup than the corresponding within-group fulfilled connection rate of that SZKB subgroup (Figure S4).

## Table legends

**Table S1. (separate file): Sample and genome data**

**Table S1a:** Sampled remains. Archaeological and storage data of the sampled skeletons based on Éry, 2008 and the handwritten report of Éry and colleagues. Radiocarbon data were generated in this study.

**Table S1s:** Detailed whole-genome sequencing (WGS) data of the analyzed individuals. Table S1c: Genetic sex and Haplogroup determination for 100 samples with low human DNA content, enriched using the myBaits Expert Human Affinities Kit - Prime Plus.

**Table S2. (separate file): PCA data**

**Table S2a:** Modern population background for the West Eurasian (EUR) PCA. The recent European individuals included in the background of Figure 2a and b.

**Table S2b:** Medieval reference samples. The European medieval individuals projected onto the PCA in Figure2b.

**Table S2c:** Modern population background for the whole Eurasian (EAS) PCA. The recent European individuals included in the background of Figure 2C.

**Table S3. (separate file): qpAdm data**

**Table S3a:** qpAdm modeling summary. Tables S3b–c show the models with significant P-values, obtained either after the base runs (when only one base-run model was significant) or after the model competition runs. In Table S3d all individuals are listed which were excluded from the model competition run with their non-passing models as well as individuals who were included in the model competition but all of their models were excluded by a putative source. For a detailed description of the analysis framework and content of specific columns see Methods section qpAdm.

**Table S3b:** qpAdm models of SZKB individuals that can be modeled using local Eur_Core populations (SZKB_LOC).

**Table S3c:** qpAdm models of SZKB individuals that can be modeled applying Asian ancestry component (SZKB_ADM).

**Table S3d:** qpAdm data of SZKB individuals for whom no valid model could be obtained, as the software excluded all sources in these cases (SZKB_NVM).

**Table S4. (separate file): IBD data**

**Table S4a:** Metadata of all individuals tested for IBD sharing with SZKB samples.

**Table S4b:** Pairwise IBD sharing data between all SZKB individuals and those listed in Table S4a.

**Table S5. (separate file): f4 statistics and hierarchical clustering data for samples lacking Asian ancestry.**

**Table S5a:** f4-statistic results of the SZKB_LOC and SZKB_NVM genomes. The reference populations used in the analysis were taken from Gerber et al. (2024, https://doi.org/10.1126/sciadv.adq5864).

**Table S5b:** Cluster assignment. Data used for assigning SZKB samples to Gerber et al. (2024) clusters. Classification was based on the distance from the centroids of the Gerber et al. (2024) clusters, as described in the Materials section.

**Table S6. (separate file): Relatedness estimates calculated by the correctKin analysis framework (Nyerki et al. 2023, https://doi.org/10.1186/s13059-023-02882-4).** For a detailed description on the content of the columns see Methods section Genetic sex and haplogroup determination, kinship.

**Table S7. (separate file): Group-based IBD sharing data**

**Table S7a:** IBD sharing data comparing the combined SZKB group with 64 medieval European and relevant Asian populations. Individuals from each group are listed in Table S4A.

**Table S7b:** IBD sharing data comparing the SZKB_ADM subgroup with 64 medieval European and relevant Asian populations, as well as with the other two SZKB subgroups. Individuals from each group are listed in Table S4A.

**Table S7c:** IBD sharing data comparing the SZKB_LOC subgroup with 64 medieval European and relevant Asian populations, as well as with the other two SZKB subgroups. Individuals from each group are listed in Table S4A.

**Table S7d:** IBD sharing data comparing the SZKB_NVM subgroup with 64 medieval European and relevant Asian populations, as well as with the other two SZKB subgroups. Individuals from each group are listed in Table S4A.

## Acknowledgments

The authors express their gratitude to Péter Erdő Cardinal, Archbishop of Esztergom-Budapest for the permission to exhume the human remains from the Matthias Church in Budapest. We are grateful to diocesan András Veres and to commissary Ferenc Reisner from the Diocese of Győr to enable the accession of the bone material. We thank Miklós Kásler for initiating the investigation. We also like to thank Gábor Horváth-Lugossy and to the leadership of the Szent István Museum of Székesfehérvár, Kovács Loránd Olivér and to mayor of Székesfehérvár Cser-Palkovics András for providing the conditions to allow acces<u>s</u> to the ossuary of the National Memorial in Székesfehérvár.

## Funding

This research was funded by grants from the National Research, Development and Innovation Office (TUDFO/5157-629 1/2019-ITM and TKP2020-NKA-23) to E.N. and grant from the Ministry of Culture and Innovation (MCI-670-19/2023/FÁFIN) to T.T. and E.N. This research was partially funded by the Competence Centre of the Life Sciences Cluster of the Centre of Excellence for Interdisciplinary Research, Development and Innovation of the University of Szeged to E.N., Z.M., and T.T. (the authors are members of the ‘Ancient and Modern Human Genomics Competence Center’ research group).

## Author contributions

Conceptualization: GIBV, EN, TT

Data curation: ZM, OS, ENY, GIBV

Formal analysis, Methodology and Validation: ZM, OS, GIBV

Funding acquisition: EN, TT, ZM

Investigation: OS, ZM, KM, LK, BK^1^ (Bence Kovács), AG, BT, OAV, PK, MD, GIBV

Project administration: GIBV

Resources: BK^2^ (Balázs Kertész), FSZ, ZSZ, PLN, JO

Software: ZM, OS, GIBV

Supervision: GIBV, TT Visualization: OS, GIBV, ZSZ

Writing – original draft: GIBV, TT

Writing – review & editing: all authors contributed.

## Competing interests

Authors declare that they have no competing interests.

## Data and materials availability

The raw nucleotide sequence data of the samples were deposited to the European Nucleotide Archive (http://www.ebi.ac.uk/ena) under accession number: ENA: PRJEB111039. All data are available in the main text or the supplementary materials.

## Notes

### Competing Interest Statement

P eter Lajos Nagy is the founder and owner of Praxis Genomics LLC

## References

1. P. Engel, Temetkezések a középkori székesfehérvári bazilikában. SZÁZADOK Magy. Tört. Társ. FOLYÓIRATA, 613–637 (1987).

2. G. Thoroczkay, “A székesfehérvári bazilika és prépostság az Árpád-korban” in Ismeretlen Árpád-Kor. Püspökök, Legendák, Krónikák (L’Harmattan, Budapest, 2016), pp. 141–183.

3. P. Lővei, “A Szűz Mária-prépostság temploma mint temetkezőhely” in Solium Regni – Az Ország Trónusa. A Koronázótemplom Története És Faragványai (Szent István Király Múzeum, Székesfehérvár, 2024), pp. 104–128.

4. T. Kádár, A középkori magyar királyi és kormányzói családok tagjainak elhalálozási és temetési időpontjai valamint helyei 1301–1541 között. Fons, 25–53 (2011).

5. B. Kertész, A. Ribi, R. Skorka, B. Weisz, A. Zsoldos, “A prépostsági templomban eltemetett főurak és prépostok életútja” in Solium Regni – Az Ország Trónusa. A Koronázótemplom Története És Faragványai (Szent István Király Múzeum, Székesfehérvár, 2024), pp. 323–329.

6. K. Éry, *A SZÉKESFEHÉRVÁRI KIRÁLYI BAZILIKA EMBERTAN LELETEI 1848-*2002 (Balassi Kiadó, Budapest, 2008).

7. I. Hankó, Királyaink Tömegsírban (Magyar Ház, 2004).

8. A. Kralovánszky, Előzetes jelentés az 1965. évi székesfehérvári feltárásokról. Alba Regia, 253–262 (1968).

9. A. Kralovánszky, “A székesfehérvári királyi bazilika alapításának kérdéséhez (Zur Frage der Gründung der königlichen Basilika von Székesfehérvár)” in A Móra Ferenc Múzeum Évkönyve 1966–67/2 (Móra Ferenc Múzeum, Szeged, 1968), pp. 121–125.

10. P. Biczó, *Solium Regni – Az Ország Trónusa*. A Koronázótemplom Története És Faragványai (Szent István Király Múzeum, Székesfehérvár, 2024).

11. Z. Szabó, *A Székesfehérvári Királyi Bazilika Építéstörténete* (Balassi Kiadó, Budapest, 2018)vol. II/2A.

12. Z. Bánki, Régészeti kutatások. Alba Regia, 179–180 (1968).

13. G. I. B. Varga, K. Maár, A. Ginguta, B. Kovács, B. Tihanyi, L. Kis, O. Váradi, P. Kiss, D. Szokolóczi, O. Schütz, Z. Maróti, E. Nyerki, I. Nagy, D. Latinovics, T. Török, E. Neparáczki, An archaeogenetic approach to identify the remains of the Hungarian Kings. Working Plan. Ephemer. Hung., 333–342 (2021).

14. B. Kovács, J. Olasz, Z. Maróti, O. Schütz, N. Rouse, M. F. Nagy, A. Ginguta, K. Maár, B. Tihanyi, L. Kis, B. Holczmann, B. Kertész, Z. Szabó, Z. Bernert, E. Neparáczki, T. Török, M. Kásler, P. L. Nagy, G. I. B. Varga, Newly discovered Árpád dynasty members from the Ossuary of the Royal Basilica at Székesfehérvár. Manuscript (2026).

15. C. E. G. Amorim, S. Vai, C. Posth, A. Modi, I. Koncz, S. Hakenbeck, M. C. La Rocca, B. Mende, D. Bobo, W. Pohl, L. P. Baricco, E. Bedini, P. Francalacci, C. Giostra, T. Vida, D. Winger, U. von Freeden, S. Ghirotto, M. Lari, G. Barbujani, J. Krause, D. Caramelli, P. J. Geary, K. R. Veeramah, Understanding 6th-century barbarian social organization and migration through paleogenomics. Nat. Commun. 9, 3547 (2018).

16. D. N. Vyas, I. Koncz, A. Modi, B. G. Mende, Y. Tian, P. Francalacci, M. Lari, S. Vai, P. Straub, Z. Gallina, T. Szeniczey, T. Hajdu, L. Pejrani Baricco, C. Giostra, R. Radzevičiūtė, Z. Hofmanová, S. Évinger, Z. Bernert, W. Pohl, D. Caramelli, T. Vida, P. J. Geary, K. R. Veeramah, Fine-scale sampling uncovers the complexity of migrations in 5th–6th century Pannonia. Curr. Biol. 33, 3951–3961.e11 (2023).

17. Z. Maróti, E. Neparáczki, O. Schütz, K. Maár, G. I. B. Varga, B. Kovács, T. Kalmár, E. Nyerki, I. Nagy, D. Latinovics, B. Tihanyi, A. Marcsik, G. Pálfi, Z. Bernert, Z. Gallina, C. Horváth, S. Varga, L. Költő, I. Raskó, P. L. Nagy, C. Balogh, A. Zink, F. Maixner, A. Götherström, R. George, C. Szalontai, G. Szenthe, E. Gáll, A. P. Kiss, B. Gulyás, B. Ny. Kovacsóczy, S. S. Gál, P. Tomka, T. Török, The genetic origin of Huns, Avars, and conquering Hungarians. Curr. Biol. 32, 2858–2870.e7 (2022).

18. B. Gyuris, L. Vyazov, A. Türk, P. Flegontov, B. Szeifert, P. Langó, B. G. Mende, V. Csáky, A. A. Chizhevskiy, I. R. Gazimzyanov, A. A. Khokhlov, A. G. Kolonskikh, N. P. Matveeva, R. R. Ruslanova, M. P. Rykun, A. Sitdikov, E. V. Volkova, S. G. Botalov, D. G. Bugrov, I. V. Grudochko, O. Komar, A. A. Krasnoperov, O. E. Poshekhonova, I. Chikunova, F. Sungatov, D. A. Stashenkov, S. Zubov, A. S. Zelenkov, H. Ringbauer, O. Cheronet, R. Pinhasi, A. Akbari, N. Rohland, S. Mallick, D. Reich, A. Szécsényi-Nagy, Long shared haplotypes identify the southern Urals as a primary source for the 10th-century Hungarians. Cell 188, 6064–6078.e11 (2025).

19. G. A. Gnecchi-Ruscone, A. Szécsényi-Nagy, I. Koncz, G. Csiky, Z. Rácz, A. B. Rohrlach, G. Brandt, N. Rohland, V. Csáky, O. Cheronet, B. Szeifert, T. Á. Rácz, A. Benedek, Z. Bernert, N. Berta, S. Czifra, J. Dani, Z. Farkas, T. Hága, T. Hajdu, M. Jászberényi, V. Kisjuhász, B. Kolozsi, P. Major, A. Marcsik, B. Ny. Kovacsóczy, C. Balogh, G. M. Lezsák, J. G. Ódor, M. Szelekovszky, T. Szeniczey, J. Tárnoki, Z. Tóth, E. K. Tutkovics, B. G. Mende, P. Geary, W. Pohl, T. Vida, R. Pinhasi, D. Reich, Z. Hofmanová, C. Jeong, J. Krause, Ancient genomes reveal origin and rapid *trans*-Eurasian migration of 7th century Avar elites. Cell 185, 1402–1413.e21 (2022).

20. D. Gerber, V. Csáky, B. Szeifert, N. Borbély, K. Jakab, G. Mező, Z. Petkes, F. Szücsi, S. Évinger, C. Líbor, P. Rácz, K. Kiss, B. G. Mende, B. M. Szőke, A. Szécsényi-Nagy, Ancient genomes reveal Avar-Hungarian transformations in the 9th-10th centuries CE Carpathian Basin. Sci. Adv. 10, eadq5864 (2024).

21. P. L. Nagy, J. Olasz, E. Neparáczki, N. Rouse, K. Kapuria, S. Cano, H. Chen, J. Di Cristofaro, G. Runfeldt, N. Ekomasova, Z. Maróti, J. Jeney, S. Litvinov, M. Dzhaubermezov, L. Gabidullina, Z. Szentirmay, G. Szabados, D. Zgonjanin, J. Chiaroni, D. M. Behar, E. Khusnutdinova, P. A. Underhill, M. Kásler, Determination of the phylogenetic origins of the Árpád Dynasty based on Y chromosome sequencing of Béla the Third. Eur. J. Hum. Genet. 29, 164–172 (2021).

22. G. Renaud, V. Slon, A. T. Duggan, J. Kelso, Schmutzi: estimation of contamination and endogenous mitochondrial consensus calling for ancient DNA. Genome Biol. 16, 224 (2015).

23. T. S. Korneliussen, A. Albrechtsen, R. Nielsen, ANGSD: Analysis of Next Generation Sequencing Data. BMC Bioinformatics 15, 356 (2014).

24. G. I. B. Varga, L. A. Kristóf, K. Maár, L. Kis, O. Schütz, O. Váradi, B. Kovács, A. Gînguță, B. Tihanyi, P. L. Nagy, Z. Maróti, E. Nyerki, T. Török, E. Neparáczki, The archaeogenomic validation of Saint Ladislaus’ relic provides insights into the Árpád dynasty’s genealogy. J. Genet. Genomics 50, 58–61 (2023).

25. A. Gînguță, B. Kovács, O. Schütz, B. Tihanyi, E. Nyerki, K. Maár, Z. Maróti, G. I. B. Varga, D. Băcueț-Crișan, T. Keresztes, T. Török, E. Neparáczki, Genetic identification of members of the prominent Báthory aristocratic family. iScience 26, 107911 (2023).

26. I. Stolarek, M. Zenczak, L. Handschuh, A. Juras, M. Marcinkowska-Swojak, A. Spinek, A. Dębski, M. Matla, H. Kóčka-Krenz, J. Piontek, M. Figlerowicz, Polish Archaeogenomics Consortium Team, Genetic history of East-Central Europe in the first millennium CE. Genome Biol. 24, 173 (2023).

27. A. Margaryan, D. J. Lawson, M. Sikora, F. Racimo, S. Rasmussen, I. Moltke, L. M. Cassidy, E. Jørsboe, A. Ingason, M. W. Pedersen, T. Korneliussen, H. Wilhelmson, M. M. Buś, P. de Barros Damgaard, R. Martiniano, G. Renaud, C. Bhérer, J. V. Moreno-Mayar, A. K. Fotakis, M. Allen, R. Allmäe, M. Molak, E. Cappellini, G. Scorrano, H. McColl, A. Buzhilova, A. Fox, A. Albrechtsen, B. Schütz, B. Skar, C. Arcini, C. Falys, C. H. Jonson, D. Błaszczyk, D. Pezhemsky, G. Turner-Walker, H. Gestsdóttir, I. Lundstrøm, I. Gustin, I. Mainland, I. Potekhina, I. M. Muntoni, J. Cheng, J. Stenderup, J. Ma, J. Gibson, J. Peets, J. Gustafsson, K. H. Iversen, L. Simpson, L. Strand, L. Loe, M. Sikora, M. Florek, M. Vretemark, M. Redknap, M. Bajka, T. Pushkina, M. Søvsø, N. Grigoreva, T. Christensen, O. Kastholm, O. Uldum, P. Favia, P. Holck, S. Sten, S. V. Arge, S. Ellingvåg, V. Moiseyev, W. Bogdanowicz, Y. Magnusson, L. Orlando, P. Pentz, M. D. Jessen, A. Pedersen, M. Collard, D. G. Bradley, M. L. Jørkov, J. Arneborg, N. Lynnerup, N. Price, M. T. P. Gilbert, M. E. Allentoft, J. Bill, S. M. Sindbæk, L. Hedeager, K. Kristiansen, R. Nielsen, T. Werge, E. Willerslev, Population genomics of the Viking world. Nature 585, 390–396 (2020).

28. S. S. Ebenesersdóttir, M. Sandoval-Velasco, E. D. Gunnarsdóttir, A. Jagadeesan, V. B. Guðmundsdóttir, E. L. Thordardóttir, M. S. Einarsdóttir, K. H. S. Moore, Á. Sigurðsson, D. N. Magnúsdóttir, H. Jónsson, S. Snorradóttir, E. Hovig, P. Møller, I. Kockum, T. Olsson, L. Alfredsson, T. F. Hansen, T. Werge, G. L. Cavalleri, E. Gilbert, C. Lalueza-Fox, J. W. Walser, S. Kristjánsdóttir, S. Gopalakrishnan, L. Árnadóttir, Ó. Þ. Magnússon, M. T. P. Gilbert, K. Stefánsson, A. Helgason, Ancient genomes from Iceland reveal the making of a human population. Science 360, 1028–1032 (2018).

29. R. Rodríguez-Varela, K. H. S. Moore, S. S. Ebenesersdóttir, G. M. Kilinc, A. Kjellström, L. Papmehl-Dufay, C. Alfsdotter, B. Berglund, L. Alrawi, N. Kashuba, V. Sobrado, V. K. Lagerholm, E. Gilbert, G. L. Cavalleri, E. Hovig, I. Kockum, T. Olsson, L. Alfredsson, T. F. Hansen, T. Werge, A. R. Munters, C. Bernhardsson, B. Skar, A. Christophersen, G. Turner-Walker, S. Gopalakrishnan, E. Daskalaki, A. Omrak, P. Pérez-Ramallo, P. Skoglund, L. Girdland-Flink, F. Gunnarsson, C. Hedenstierna-Jonson, M. T. P. Gilbert, K. Lidén, M. Jakobsson, L. Einarsson, H. Victor, M. Krzewińska, T. Zachrisson, J. Storå, K. Stefánsson, A. Helgason, A. Götherström, The genetic history of Scandinavia from the Roman Iron Age to the present. Cell 186, 32–46.e19 (2023).

30. I. Olalde, P. Carrión, I. Mikić, N. Rohland, S. Mallick, I. Lazaridis, M. Mah, M. Korać, S. Golubović, S. Petković, N. Miladinović-Radmilović, D. Vulović, T. Alihodžić, A. Ash, M. Baeta, J. Bartík, Ž. Bedić, M. Bilić, C. Bonsall, M. Bunčić, D. Bužanić, M. Carić, L. Čataj, M. Cvetko, I. Drnić, A. Dugonjić, A. Đukić, K. Đukić, Z. Farkaš, P. Jelínek, M. Jovanovic, I. Kaić, H. Kalafatić, M. Krmpotić, S. Krznar, T. Leleković, M. M. de Pancorbo, V. Matijević, B. Milošević Zakić, A. J. Osterholtz, J. M. Paige, D. Tresić Pavičić, Z. Premužić, P. Rajić Šikanjić, A. Rapan Papeša, L. Paraman, M. Sanader, I. Radovanović, M. Roksandic, A. Šefčáková, S. Stefanović, M. Teschler-Nicola, D. Tončinić, B. Zagorc, K. Callan, F. Candilio, O. Cheronet, D. Fernandes, A. Kearns, A. M. Lawson, K. Mandl, A. Wagner, F. Zalzala, A. Zettl, Ž. Tomanović, D. Keckarević, M. Novak, K. Harper, M. McCormick, R. Pinhasi, M. Grbić, C. Lalueza-Fox, D. Reich, A genetic history of the Balkans from Roman frontier to Slavic migrations. Cell 186, 5472–5485.e9 (2023).

31. S. Aneli, T. Saupe, F. Montinaro, A. Solnik, L. Molinaro, C. Scaggion, N. Carrara, A. Raveane, T. Kivisild, M. Metspalu, C. L. Scheib, L. Pagani, The Genetic Origin of Daunians and the Pan-Mediterranean Southern Italian Iron Age Context. Mol. Biol. Evol. 39, msac014 (2022).

32. M. L. Antonio, C. L. Weiß, Z. Gao, S. Sawyer, V. Oberreiter, H. M. Moots, J. P. Spence, O. Cheronet, B. Zagorc, E. Praxmarer, K. T. Özdoğan, L. Demetz, P. Gelabert, D. Fernandes, M. Lucci, T. Alihodžić, S. Amrani, P. Avetisyan, C. Baillif-Ducros, Ž. Bedić, A. Bertrand, M. Bilić, L. Bondioli, P. Borówka, E. Botte, J. Burmaz, D. Bužanić, F. Candilio, M. Cvetko, D. De Angelis, I. Drnić, K. Elschek, M. Fantar, A. Gaspari, G. Gasperetti, F. Genchi, S. Golubović, Z. Hukeľová, R. Jankauskas, K. J. Vučković, G. Jeremić, I. Kaić, K. Kazek, H. Khachatryan, A. Khudaverdyan, S. Kirchengast, M. Korać, V. Kozlowski, M. Krošláková, D. Kušan Špalj, F. La Pastina, M. Laguardia, S. Legrand, T. Leleković, T. Leskovar, W. Lorkiewicz, D. Los, A. M. Silva, R. Masaryk, V. Matijević, Y. M. S. Cherifi, N. Meyer, I. Mikić, N. Miladinović-Radmilović, B. Milošević Zakić, L. Nacouzi, M. Natuniewicz-Sekuła, A. Nava, C. Neugebauer-Maresch, J. Nováček, A. Osterholtz, J. Paige, L. Paraman, D. Pieri, K. Pieta, S. Pop-Lazić, M. Ruttkay, M. Sanader, A. Sołtysiak, A. Sperduti, T. Stankovic Pesterac, M. Teschler-Nicola, I. Teul, D. Tončinić, J. Trapp, D. Vulović, T. Waliszewski, D. Walter, M. Živanović, M. el M. Filah, M. Čaušević-Bully, M. Šlaus, D. Borić, M. Novak, A. Coppa, R. Pinhasi, J. K. Pritchard, Stable population structure in Europe since the Iron Age, despite high mobility. eLife 13, e79714 (2024).

33. É. Harney, N. Patterson, D. Reich, J. Wakeley, Assessing the performance of qpAdm: a statistical tool for studying population admixture. Genetics 217, iyaa045 (2021).

34. P. de B. Damgaard, N. Marchi, S. Rasmussen, M. Peyrot, G. Renaud, T. Korneliussen, J. V. Moreno-Mayar, M. W. Pedersen, A. Goldberg, E. Usmanova, N. Baimukhanov, V. Loman, L. Hedeager, A. G. Pedersen, K. Nielsen, G. Afanasiev, K. Akmatov, A. Aldashev, A. Alpaslan, G. Baimbetov, V. I. Bazaliiskii, A. Beisenov, B. Boldbaatar, B. Boldgiv, C. Dorzhu, S. Ellingvag, D. Erdenebaatar, R. Dajani, E. Dmitriev, V. Evdokimov, K. M. Frei, A. Gromov, A. Goryachev, H. Hakonarson, T. Hegay, Z. Khachatryan, R. Khaskhanov, E. Kitov, A. Kolbina, T. Kubatbek, A. Kukushkin, I. Kukushkin, N. Lau, A. Margaryan, I. Merkyte, I. V. Mertz, V. K. Mertz, E. Mijiddorj, V. Moiyesev, G. Mukhtarova, B. Nurmukhanbetov, Z. Orozbekova, I. Panyushkina, K. Pieta, V. Smrčka, I. Shevnina, A. Logvin, K.-G. Sjögren, T. Štolcová, A. M. Taravella, K. Tashbaeva, A. Tkachev, T. Tulegenov, D. Voyakin, L. Yepiskoposyan, S. Undrakhbold, V. Varfolomeev, A. Weber, M. A. Wilson Sayres, N. Kradin, M. E. Allentoft, L. Orlando, R. Nielsen, M. Sikora, E. Heyer, K. Kristiansen, E. Willerslev, 137 ancient human genomes from across the Eurasian steppes. Nature 557, 369–374 (2018).

35. L. Saag, O. Utevska, S. Zadnikov, I. Shramko, K. Gorbenko, M. Bandrivskyi, D. Pavliv, I. Bruyako, D. Grechko, V. Okatenko, G. Toshev, S. Andrukh, V. Radziyevska, Y. Buynov, V. Kotenko, O. Smyrnov, O. Petrauskas, B. Magomedov, S. Didenko, A. Heiko, R. Reida, S. Sapiehin, V. Aksonov, O. Laptiev, S. Terskyi, V. Skorokhod, V. Zhyhola, Y. Sytyi, M. Järve, C. L. Scheib, K. Anastasiadou, M. Kelly, M. Williams, M. Silva, C. Barrington, A. Gilardet, R. Macleod, P. Skoglund, M. G. Thomas, North Pontic crossroads: Mobility in Ukraine from the Bronze Age to the early modern period. Sci. Adv. 11, eadr0695 (2025).

36. E. Nyerki, T. Kalmár, O. Schütz, R. M. Lima, E. Neparáczki, T. Török, Z. Maróti, correctKin: an optimized method to infer relatedness up to the 4th degree from low-coverage ancient human genomes. Genome Biol. 24, 38 (2023).

37. H. Ringbauer, Y. Huang, A. Akbari, S. Mallick, I. Olalde, N. Patterson, D. Reich, Accurate detection of identity-by-descent segments in human ancient DNA. Nat. Genet. 56, 143–151 (2024).

38. O. Schütz, Z. Maróti, B. Tihanyi, A. P. Kiss, E. Nyerki, A. Gînguță, P. Kiss, G. I. B. Varga, B. Kovács, K. Maár, B. Ny. Kovacsóczy, N. Lukács, I. Major, A. Marcsik, E. Patyi, A. Szigeti, Z. Tóth, D. Walter, G. Wilhelm, R. Cs. Andrási, Z. Bernert, L. Kis, L. Oța, G. Pálfi, G. Pintye, D. Pópity, A. Simalcsik, A. D. Soficaru, O. Spekker, S. Varga, E. Neparáczki, T. Török, Unveiling the origins and genetic makeup of the “forgotten people”: A study of the Sarmatian-period population in the Carpathian Basin. Cell 188, 4074–4090.e11 (2025).

39. G. I. B. Varga, Z. Maróti, O. Schütz, K. Maár, E. Nyerki, B. Tihanyi, O. A. Váradi, A. Gînguță, B. Kovács, P. Kiss, M. Dosztig, Z. Gallina, T. Török, J. B. Szabó, M. Makoldi, E. Neparáczki, Archaeogenetic analysis revealed East Eurasian paternal origin to the Aba royal family of Hungary. iScience 27, 110892 (2024).

40. G. Hellenthal, G. B. J. Busby, G. Band, J. F. Wilson, C. Capelli, D. Falush, S. Myers, A Genetic Atlas of Human Admixture History. Science 343, 747–751 (2014).

41. A. Zsoldos, *The Árpáds and Their People. An Introduction to the History of Hungary from Cca*. 900 to 1301 (Research Centre for the Humanities, Budapest, 2020).

42. A. Kubinyi, “németek” in *Korai Magyar Történeti Lexikon* *(9–14. Század)* (Akadémiai Kiadó, Budapest, 1994), p. 485.

43. H. Zimmermann, “Die deutsch-ungarischen Beziehungen in der Mitte des 12. Jahrhunderts und die Berufung der Siebenbürger Sachsen” in Siebenbürgen Und Seine Hospites Theutonici. Vorträge Und Forschungen Zur Südostdeutschen Geschichte. Festgabe Zum 70. Geburtstag von Harald Zimmermann (Böhlau Köln, Köln, 1996), pp. 83–101.

44. P. Lökös, V. Paál, “németek” in Magyar Művelődéstörténeti Lexikon, Középkor És Kora Újkor. I–XIII. (Balassi Kiadó, Budapest, 2003).

45. M. Molnár, R. Janovics, I. Major, J. Orsovszki, R. Gönczi, M. Veres, A. G. Leonard, S. M. Castle, T. E. Lange, L. Wacker, I. Hajdas, A. J. T. Jull, Status Report of the New AMS 14C Sample Preparation Lab of the Hertelendi Laboratory of Environmental Studies (Debrecen, Hungary). Radiocarbon 55, 665–676 (2013).

46. P. J. Reimer, W. E. N. Austin, E. Bard, A. Bayliss, P. G. Blackwell, C. B. Ramsey, M. Butzin, H. Cheng, R. L. Edwards, M. Friedrich, P. M. Grootes, T. P. Guilderson, I. Hajdas, T. J. Heaton, A. G. Hogg, K. A. Hughen, B. Kromer, S. W. Manning, R. Muscheler, J. G. Palmer, C. Pearson, J. van der Plicht, R. W. Reimer, D. A. Richards, E. M. Scott, J. R. Southon, C. S. M. Turney, L. Wacker, F. Adolphi, U. Büntgen, M. Capano, S. M. Fahrni, A. Fogtmann-Schulz, R. Friedrich, P. Köhler, S. Kudsk, F. Miyake, J. Olsen, F. Reinig, M. Sakamoto, A. Sookdeo, S. Talamo, The IntCal20 Northern Hemisphere Radiocarbon Age Calibration Curve (0–55 cal kBP). Radiocarbon 62, 725–757 (2020).

47. C. B. Ramsey, Bayesian Analysis of Radiocarbon Dates. Radiocarbon 51, 337–360 (2009).

48. C. B. Ramsey, Methods for Summarizing Radiocarbon Datasets. Radiocarbon 59, 1809–1833 (2017).

49. C. B. Ramsey, OxCal, version 4.4.4 (2021); https://c14.arch.ox.ac.uk/oxcal.html.

50. M. Martin, Cutadapt removes adapter sequences from high-throughput sequencing reads. EMBnet.journal 17, 10–12 (2011).

51. S. Andrews, FastQC. A quality control tool for high throughput sequence data, (2016); https://www.bioinformatics.babraham.ac.uk/projects/fastqc/.

52. H. Li, R. Durbin, Fast and accurate short read alignment with Burrows–Wheeler transform. Bioinformatics 25, 1754–1760 (2009).

53. H. Li, B. Handsaker, A. Wysoker, T. Fennell, J. Ruan, N. Homer, G. Marth, G. Abecasis, R. Durbin, 1000 Genome Project Data Processing Subgroup, The Sequence Alignment/Map format and SAMtools. Bioinformatics 25, 2078–2079 (2009).

54. Broad Institute, Picard tools, (2016); https://broadinstitute.github.io/picard/.

55. V. Link, A. Kousathanas, K. Veeramah, C. Sell, A. Scheu, D. Wegmann, ATLAS: Analysis Tools for Low-depth and Ancient Samples. bioRxiv [Preprint] (2017). 10.1101/105346.

56. M. Rasmussen, X. Guo, Y. Wang, K. E. Lohmueller, S. Rasmussen, A. Albrechtsen, L. Skotte, S. Lindgreen, M. Metspalu, T. Jombart, T. Kivisild, W. Zhai, A. Eriksson, A. Manica, L. Orlando, F. M. De La Vega, S. Tridico, E. Metspalu, K. Nielsen, M. C. Ávila-Arcos, J. V. Moreno-Mayar, C. Muller, J. Dortch, M. T. P. Gilbert, O. Lund, A. Wesolowska, M. Karmin, L. A. Weinert, B. Wang, J. Li, S. Tai, F. Xiao, T. Hanihara, G. van Driem, A. R. Jha, F.-X. Ricaut, P. de Knijf, A. B. Migliano, I. Gallego Romero, K. Kristiansen, D. M. Lambert, S. Brunak, P. Forster, B. Brinkmann, O. Nehlich, M. Bunce, M. Richards, R. Gupta, C. D. Bustamante, A. Krogh, R. A. Foley, M. M. Lahr, F. Balloux, T. Sicheritz-Pontén, R. Villems, R. Nielsen, J. Wang, E. Willerslev, An Aboriginal Australian Genome Reveals Separate Human Dispersals into Asia. Science 334, 94–98 (2011).

57. P. Skoglund, J. Storå, A. Götherström, M. Jakobsson, Accurate sex identification of ancient human remains using DNA shotgun sequencing. J. Archaeol. Sci. 40, 4477–4482 (2013).

58. B. S. Pedersen, A. R. Quinlan, Mosdepth: quick coverage calculation for genomes and exomes. Bioinformatics 34, 867–868 (2018).

59. H. Weissensteiner, D. Pacher, A. Kloss-Brandstätter, L. Forer, G. Specht, H.-J. Bandelt, F. Kronenberg, A. Salas, S. Schönherr, HaploGrep 2: mitochondrial haplogroup classification in the era of high-throughput sequencing. Nucleic Acids Res. 44, W58–W63 (2016).

60. A. Ralf, D. Montiel González, K. Zhong, M. Kayser, Yleaf: Software for Human Y-Chromosomal Haplogroup Inference from Next-Generation Sequencing Data. Mol. Biol. Evol. 35, 1291–1294 (2018).

61. International Society of Genetic Genealogy. Y-DNA Haplogroup Tree 2019-2020, version 15.73 (2020); http://www.isogg.org/tree/.

62. E. Neparáczki, L. Kis, Z. Maróti, B. Kovács, G. I. B. Varga, M. Makoldi, P. Horolma, Éva Teiszler, B. Tihanyi, P. L. Nagy, K. Maár, A. Gyenesei, O. Schütz, E. Dudás, T. Török, V. Pascuttini-Juraga, I. Peharda, L. T. Vizi, G. Horváth-Lugossy, M. Kásler, The genetic legacy of the Hunyadi descendants. Heliyon 8, e11731 (2022).

63. N. Patterson, A. L. Price, D. Reich, Population Structure and Eigenanalysis. PLOS Genet. 2, e190 (2006).

64. N. Patterson, P. Moorjani, Y. Luo, S. Mallick, N. Rohland, Y. Zhan, T. Genschoreck, T. Webster, D. Reich, Ancient Admixture in Human History. Genetics 192, 1065–1093 (2012).

65. H. Wickham, Ggplot2 (Springer International Publishing, Cham, 2016; http://link.springer.com/10.1007/978-3-319-24277-4)*Use R!*

66. W. N. Venables, B. D. Ripley, *Modern Applied Statistics with S* (Springer Nature, New York, 2002)Statistics and Computing.

67. H. Wickham, R. Francois, L. Henry, K. Müller, D. Vaughan, dplyr: A Grammar of Data Manipulation, (2026); https://dplyr.tidyverse.org.

68. H. Wickham, D. Vaughan, M. Girlich, tidyr: Tidy Messy Data, (2026); https://tidyr.tidyverse.org.

69. C. Sievert, *Interactive Web-Based Data Visualization with R, Plotly, and Shiny* (Chapman and Hall/CRC, 2020; https://plotly-r.com).

70. S. Rubinacci, D. M. Ribeiro, R. J. Hofmeister, O. Delaneau, Eficient phasing and imputation of low-coverage sequencing data using large reference panels. Nat. Genet. 53, 120–126 (2021).

71. Z. Maróti, TRUTH - True Ratio of Unbiased IBD Through Hypothesis, a tool to filter raw ancIBD segments based on experimental IBD density; https://github.com/zmaroti/ancIBDfiltration.

72. G. Csardi, T. Nepusz, The igraph software package for complex network research. InterJournal **Complex Systems**, 1695 (2006).

73. G. Csárdi, T. Nepusz, V. Traag, S. Horvát, F. Zanini, D. Noom, K. Müller, D. Schoch, M. Salmon, {igraph}: Network Analysis and Visualization in R, (2026); 10.5281/zenodo.7682609.

