## Supplementary figures for "Genomic landscape of the medieval Hungarian elite from the Székesfehérvár royal necropolis"

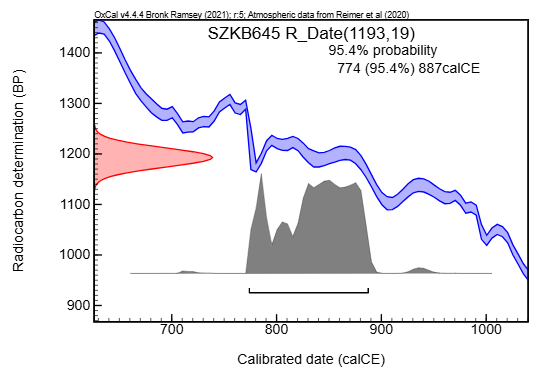

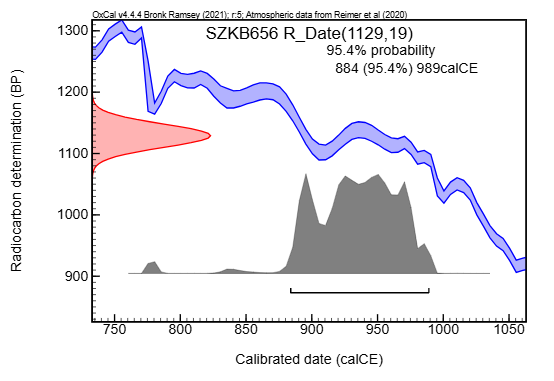

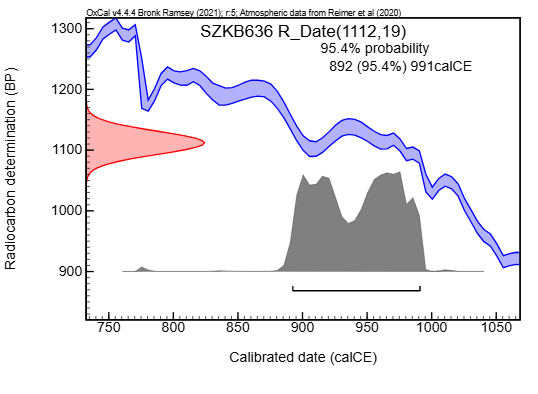

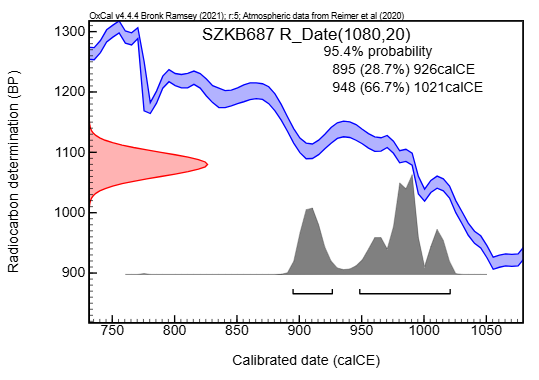

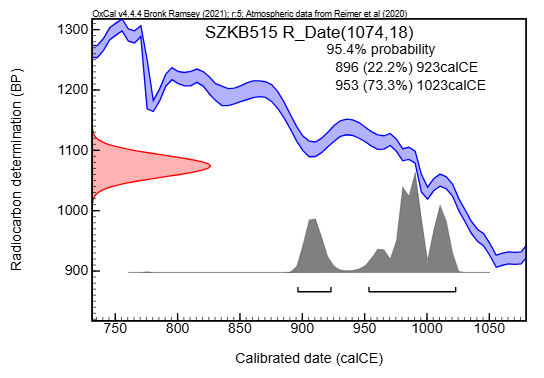

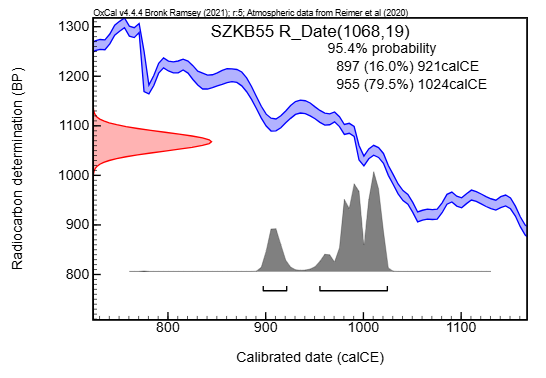

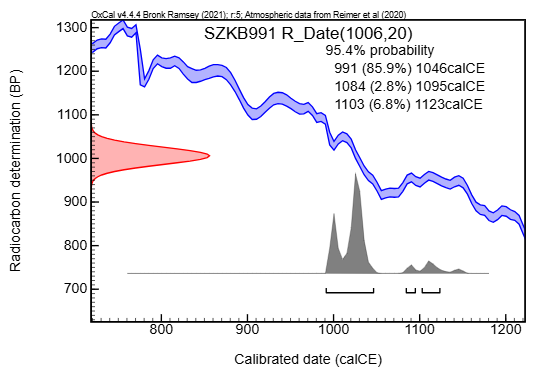

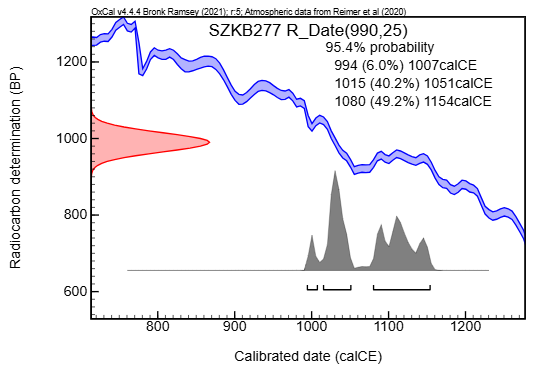

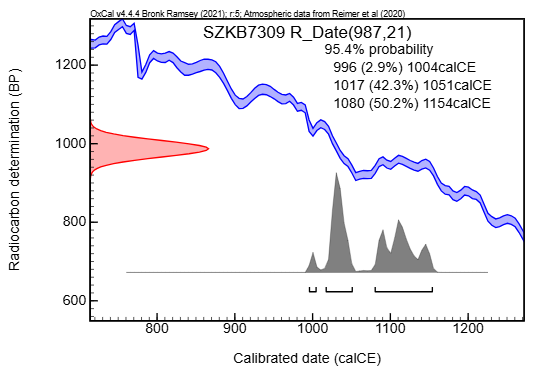

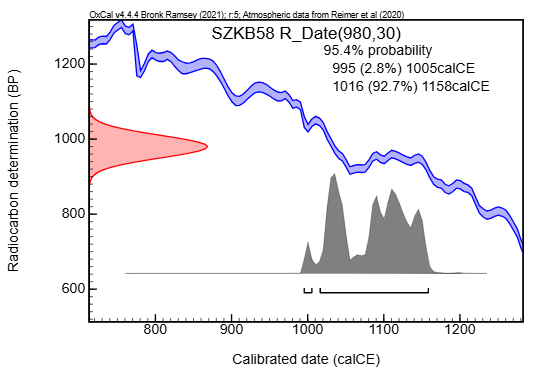

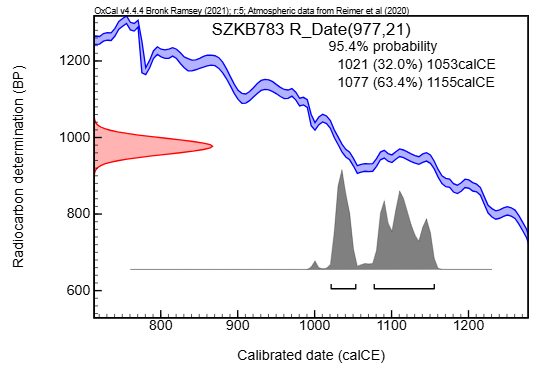

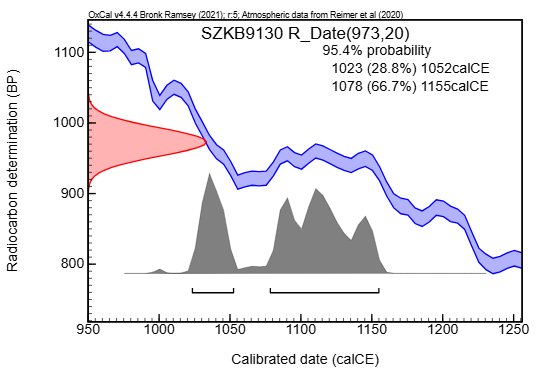

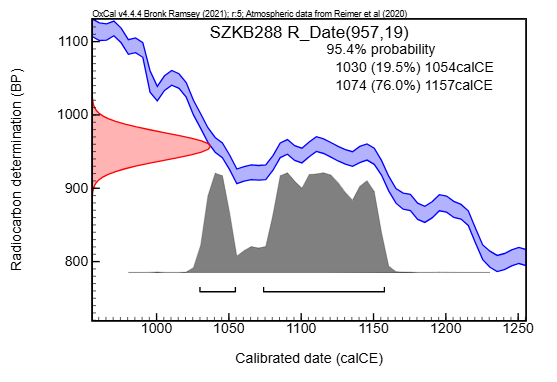

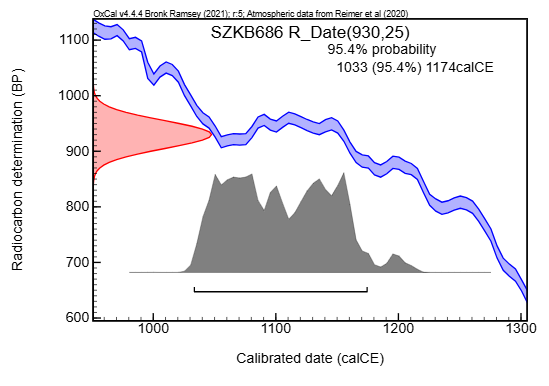

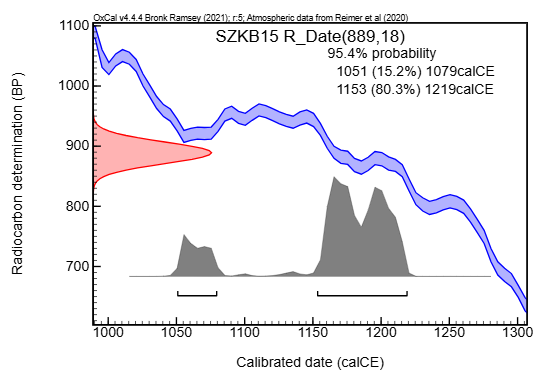

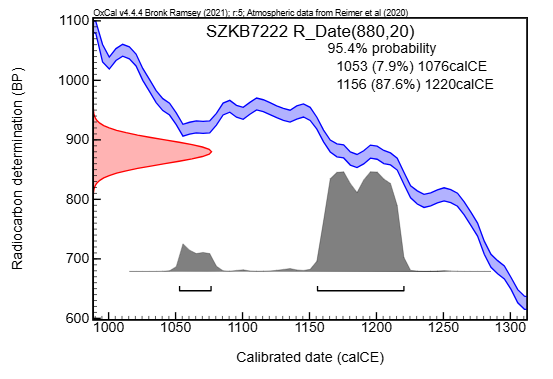

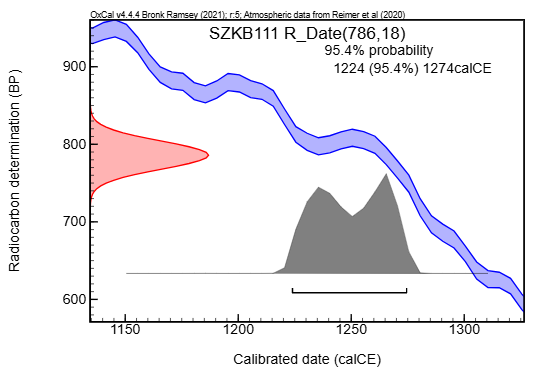

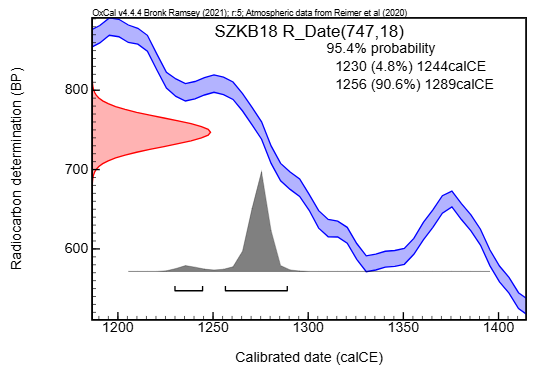

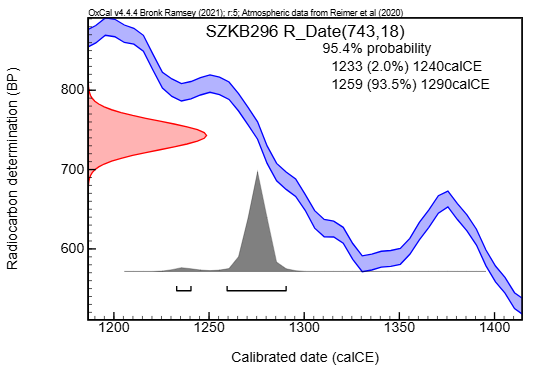

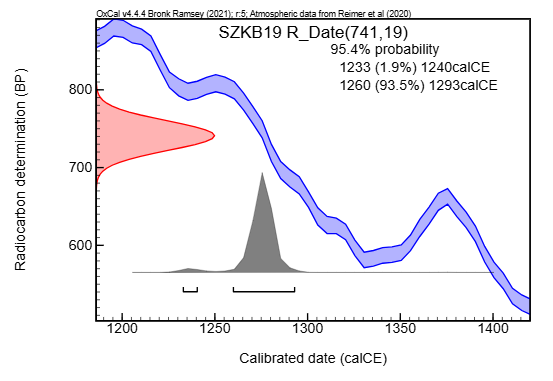

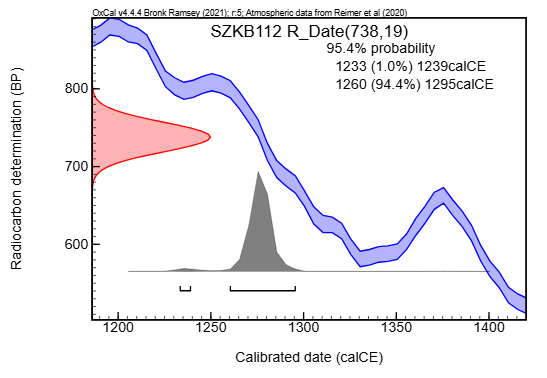

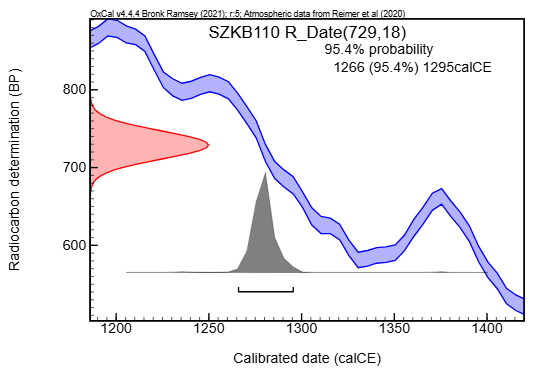

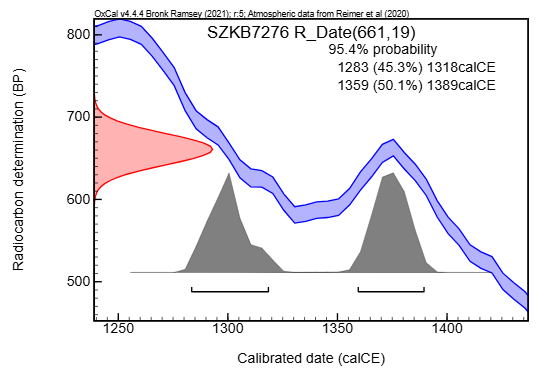

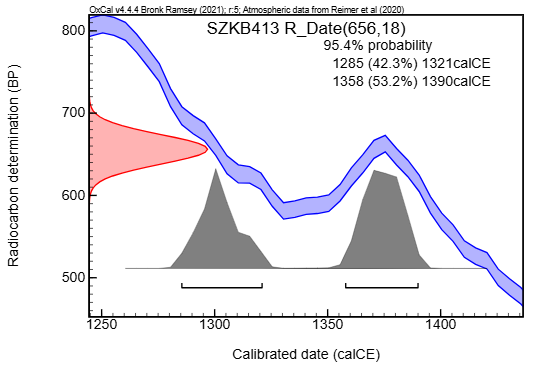

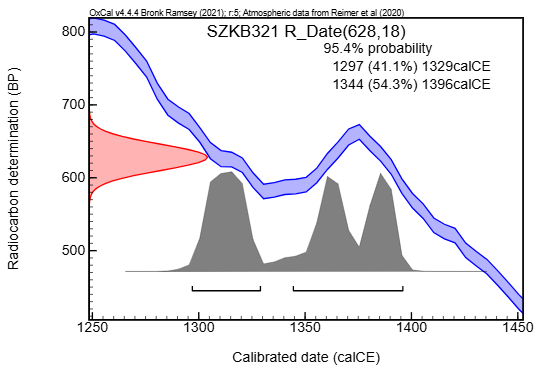

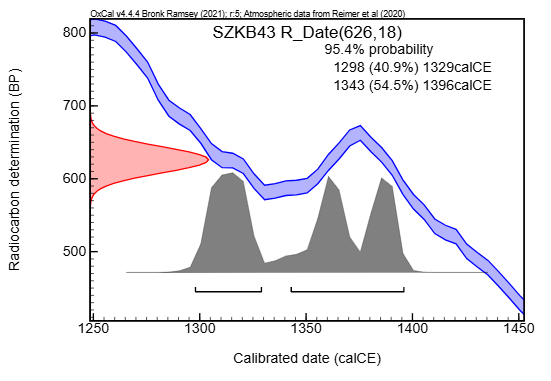

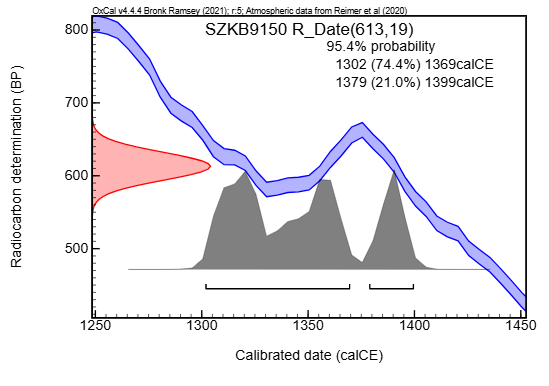

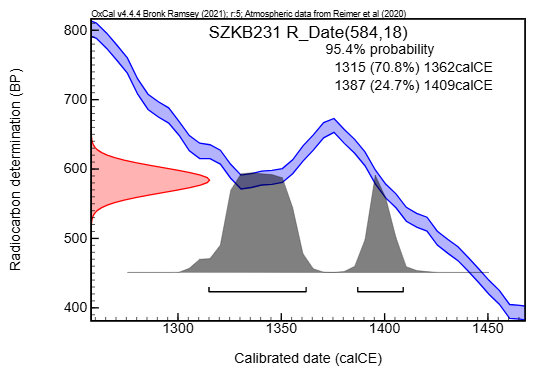

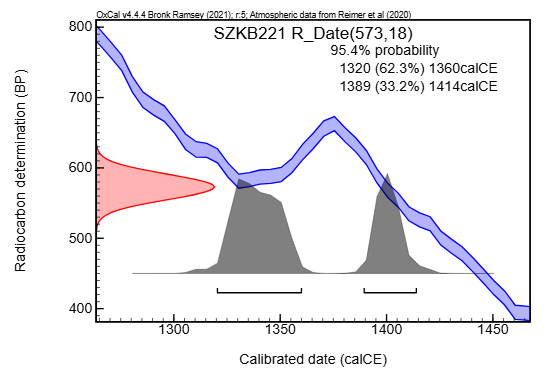

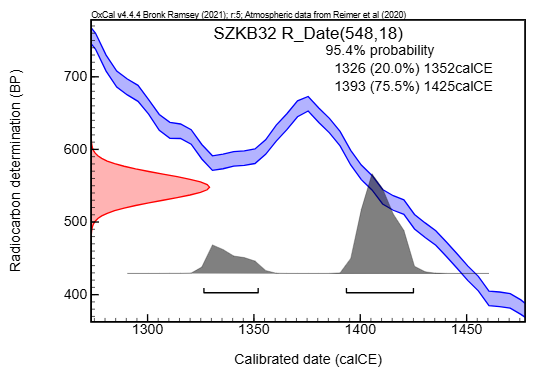

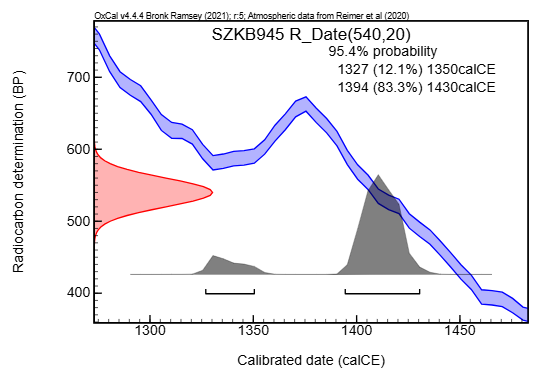

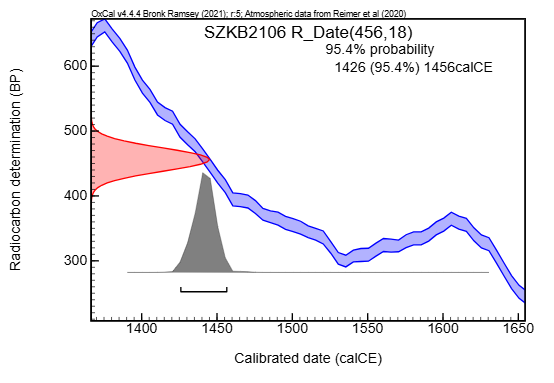

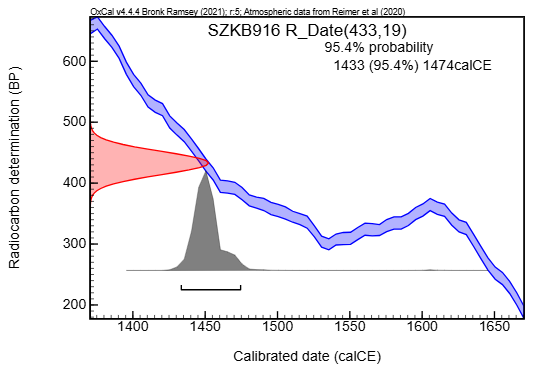

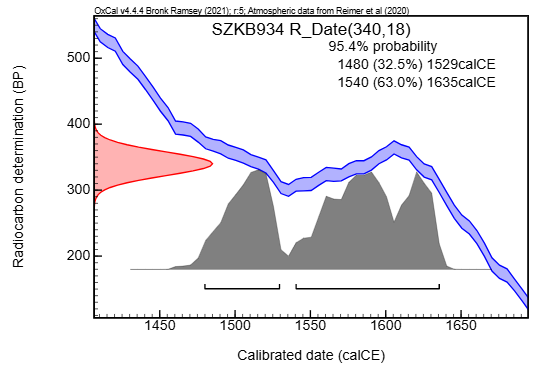

Fig. S1.

Radiocarbon calibration curves of the SZKB remains, which are summarized in Figure 1. Single plots were created by the OxCal 4.4.4 software.

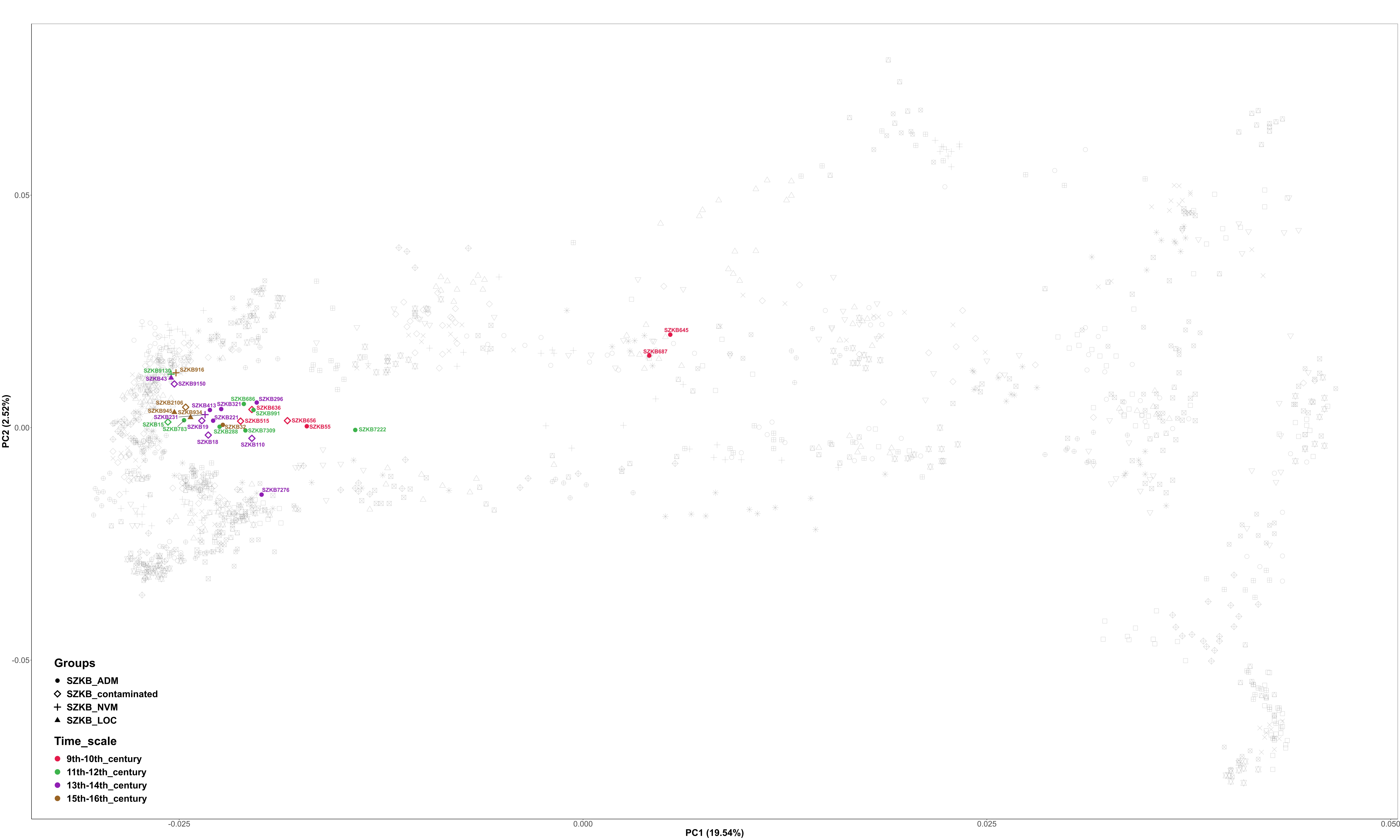

Fig. S2.

Radiocarbon dated SZKB genomes projected onto a background of present-day Eurasian populations (Maróti, 2022). The colors indicate the ages of the individuals, while the shapes of the symbols represent their SZKB subgroup affiliations, as indicated in the figure legends. A clear trend is visible in the figure: the younger the sample, the more it is shifted toward the Asian direction, reflecting an increased Asian ancestry component.

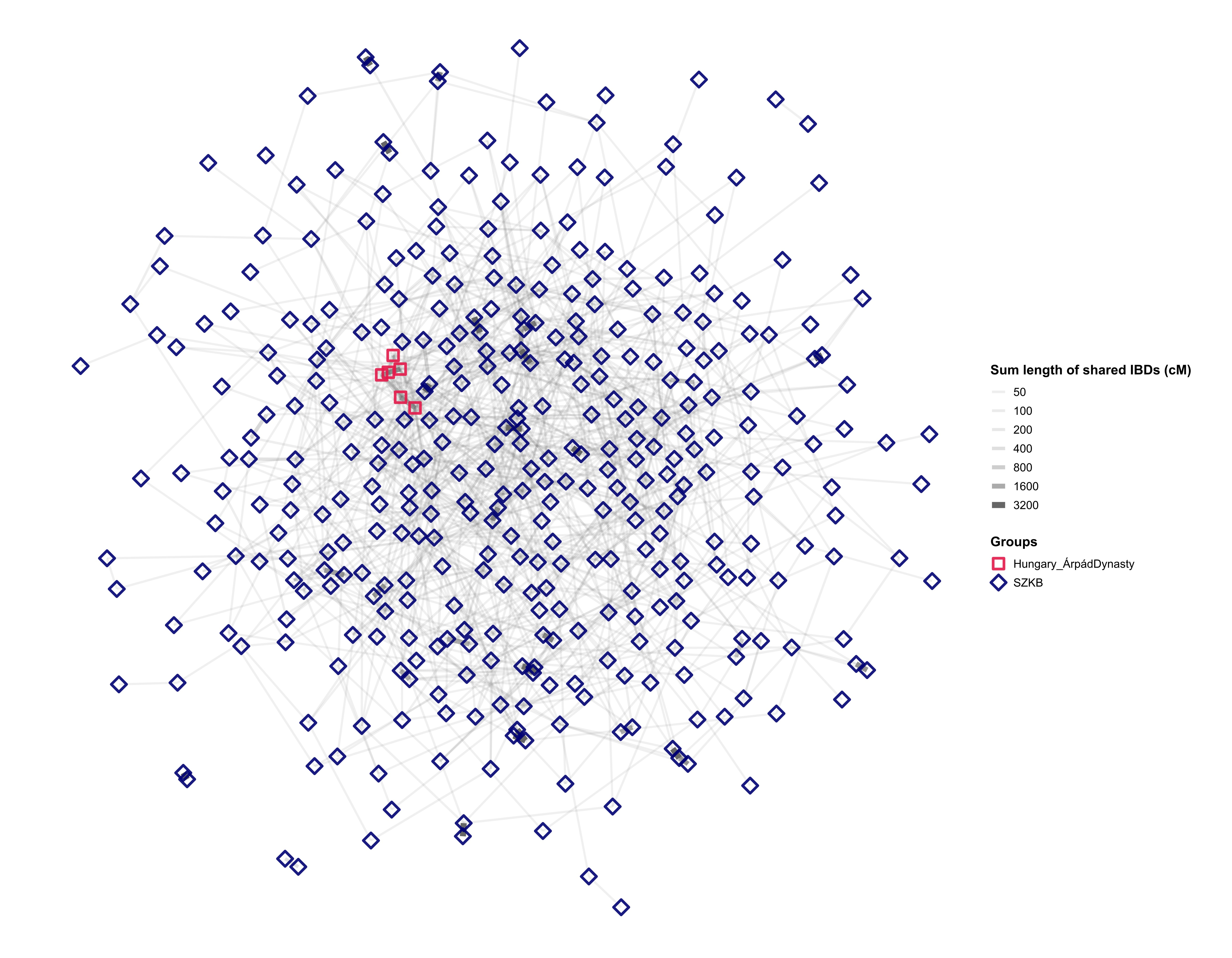

Fig. S3.

Internal IBD network of the necropolis. As most connections are based on short IBD segments, the network captures diffuse population-level structure rather than close genealogical relationships.

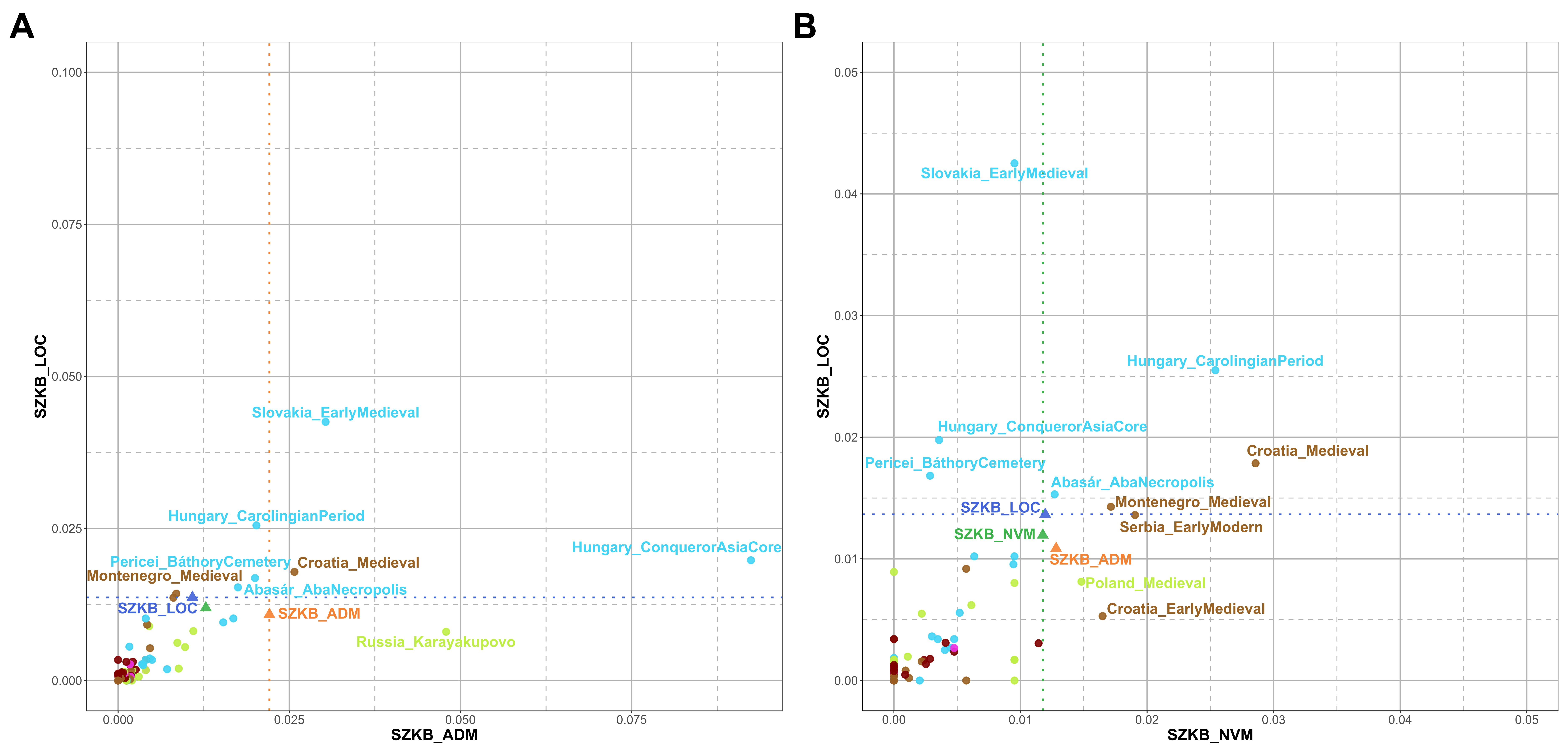

Fig. S4.

**External IBD connections of the SZKB subgroups.** **A)** External IBD connections of the SZKB_ADM subgroup, which carries Asian ancestry, and the SZKB_LOC subgroup, which can be modeled exclusively with Eur_Core. **B)** External IBD connections of the SZKB_LOC subgroup and the SZKB_NVM subgroup, for which qpAdm does not yield valid models.

The figure is based on the data presented in Table S7. We normalized the degree centrality data by dividing the detected connections by the total number of possible connections, resulting in the ratio of fulfilled connections as described in the Methods section. The values on the X and Y axes represent the ratio of fulfilled connections. The dashed line indicates the internal connectedness ratio of the given SZKB subgroup. In Figure 6, we display only those populations with the strongest connections, i.e., those falling above the dashed line.

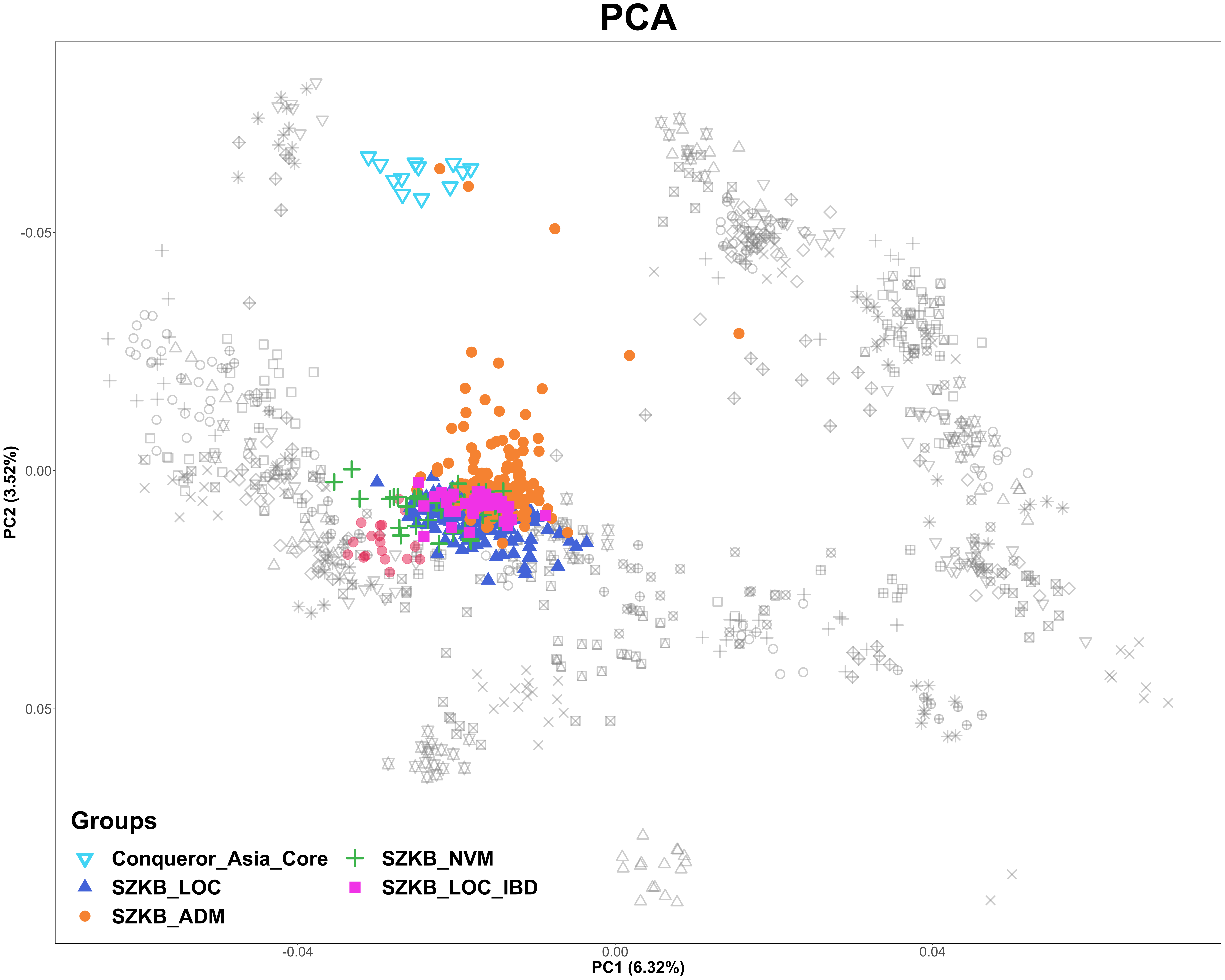

Fig. S5.

SZKB_LOC individuals exhibiting IBD sharing with Conq_Asia_Core. This figure is identical to Figure 2A, except that SZKB_LOC individuals showing IBD sharing with Conq_Asia_Core are recolored from blue triangles to magenta circles. Notably, these individuals fall precisely within the overlap zone between the SZKB_LOC and SZKB_ADM clusters on the PCA. SZKB genomes are projected onto a reference panel of present-day European individuals (14), with modern Hungarians highlighted by red dots. Conq_Asia_Core genomes (from (5)) are indicated by light blue inverted triangles. Colors denote groups based on qpAdm modeling: dark blue triangles and magenta circles represent SZKB_LOC genomes modeled from local sources; orange circles indicate SZKB_ADM genomes with significant eastern components; and green plus signs mark SZKB_NVM genomes without a valid qpAdm model.

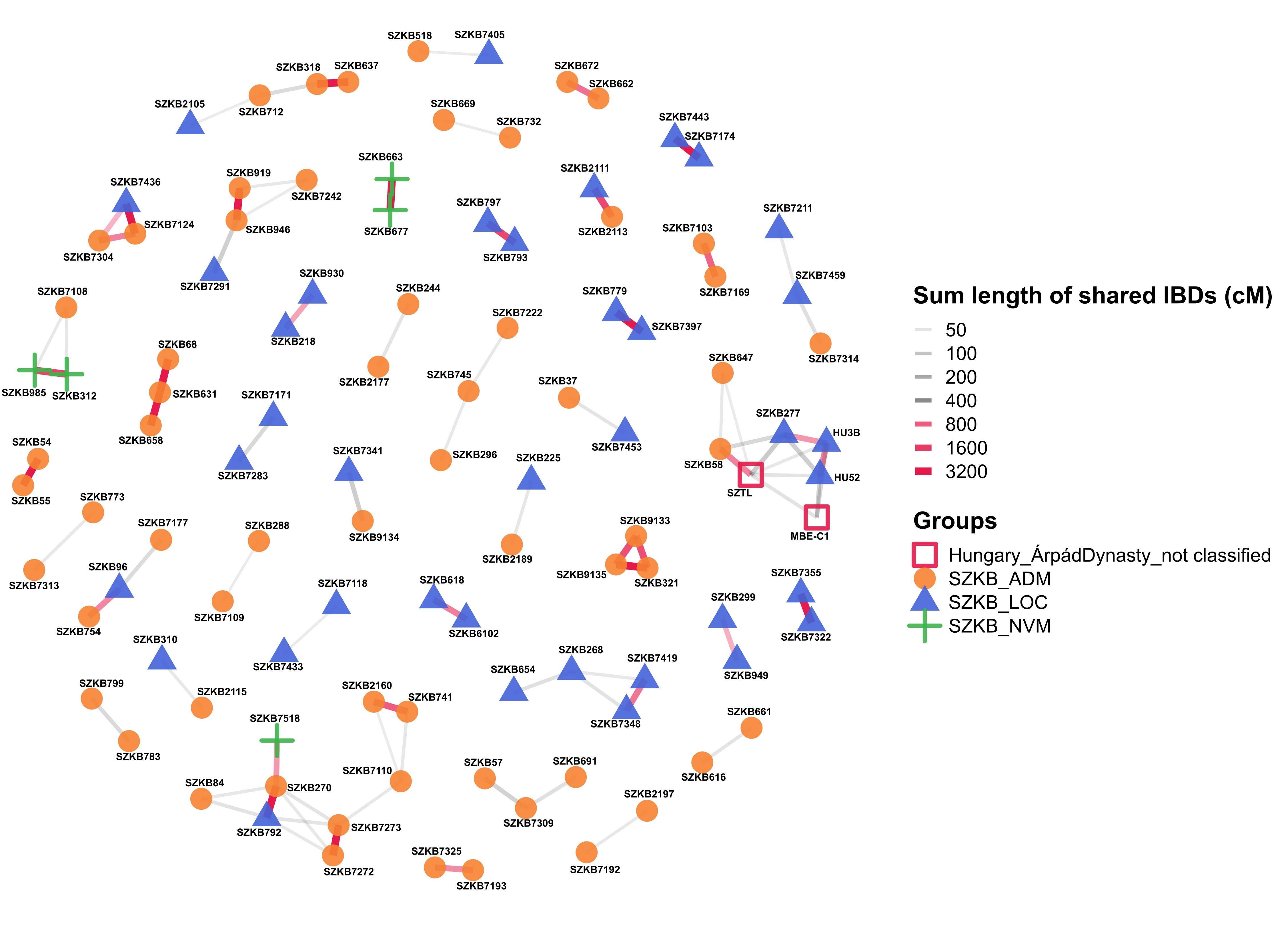

Fig. S6.

**IBD network of close genealogical connections within the necropolis. including sample IDs**

This figure is identical to Figure 5, except that sample identifiers are also displayed, and the color scheme corresponds to the SZKB subgroups: SZKB_ADM (orange circles), SZKB_LOC (blue triangles), SZKB_NVM (green plus signs), and Árpád dynasty individuals from outside Székesfehérvár (red squares).
